# Fitness selection of hyperfusogenic measles virus F proteins associated with neuropathogenic phenotypes

**DOI:** 10.1101/2020.12.22.423954

**Authors:** Satoshi Ikegame, Takao Hashiguchi, Chuan-Tien Hung, Kristina Dobrindt, Kristen J Brennand, Makoto Takeda, Benhur Lee

## Abstract

Measles virus (MeV) is resurgent and caused >200,000 deaths in 2019. MeV infection can establish a chronic latent infection of the brain that can recrudesce months to years after recovery from the primary infection. Recrudescent MeV leads to fatal subacute sclerosing panencephalitis (SSPE) or measles inclusion body encephalitis (MIBE) as the virus spreads across multiple brain regions. Most clinical isolates of SSPE/MIBE strains show mutations in the fusion (F) gene that result in a hyperfusogenic phenotype *in vitro* and allow for efficient spread in primary human neurons. Wild-type MeV receptor binding protein (RBP) is indispensable for manifesting these mutant F phenotypes, even though neurons lack canonical MeV receptors (CD150/SLAMF1 or Nectin-4). How such hyperfusogenic F mutants are selected for, and whether they confer a fitness advantage for efficient neuronal spread is unresolved. To better understand the fitness landscape that allows for the selection of such hyperfusogenic F mutants, we conducted a screen of ≥3.1×10^5^ MeV-F point mutants in their genomic context. We rescued and amplified our genomic MeV-F mutant libraries in BSR-T7 cells under conditions where MeV-F-T461I (a known SSPE mutant), but not wild-type MeV can spread. We recovered known SSPE mutants but also characterized at least 15 novel hyperfusogenic F mutations with a SSPE phenotype. Structural mapping of these mutants onto the pre-fusion MeV-F trimer confirm and extend our understanding of the fusion regulatory domains in MeV-F. Our list of hyperfusogenic F mutants is a valuable resource for future studies into MeV neuropathogenesis and the regulation of paramyxovirus fusion.

**Significance:** Measles remains a major cause of infant death globally. On rare occasions, measles virus infection of the central nervous system (CNS) leads to a fatal progressive inflammation of the brain many years after the initial infection. MeV isolates from such CNS infections harbor fusion (F) protein mutations that result in a hyperfusogenic phenotype. The small number of hyperfusogenic MeV-F mutants identified thus far limits our ability to understand how these mutations are selected in the context of CNS infections. We performed a saturating mutagenesis screen of MeV-F to identify a large set of mutants that would mimic the hyperfusogenic phenotype of MeV-F in CNS infection. Characterization of these mutants shed light on other paramyxoviruses known to establish chronic CNS infections.

## Introduction

Measles is a highly contagious acute infectious disease caused by measles virus (MeV) (Genus *morbillivirus*, Family *Paramyxoviridae*, Order *Mononegavirales*^1^). There has been a resurgence of measles in recent years due to the lack or lapse of comprehensive vaccine coverage. The global incidence of measles in 2019 of 120 per million represents a 6.7-fold increase from its nadir in 2016 (18 per million). Primary MeV infections also caused an estimated 207,500 deaths globally the same year^2^. These deaths occurred mostly in children under 5 years of age, who are also most susceptible to complications of pneumonia, or diarrhea and dehydration. Measles continue to exert its toll after recovery from acute infection. Due to virus-induced depletion of B-cell memory pools— a form of immunological amnesia— recovered children can become newly susceptible to common childhood infectious diseases ^3–5^. In the longer term, MeV can also cause chronic latent central nervous system (CNS) infections such as measles inclusion body encephalitis (MIBE) and subacute sclerosing panencephalitis (SSPE) ^6^. MIBE is restricted to patients who are immunocompromised whereas SSPE can occur in fully immunocompetent people 7-10 years after primary MeV infection^7^. The incidence of SSPE is rare; although more recent estimates of its occurrence range from 22/100,000 to 30-59/100,000 in children that acquire measles before the age of 5 ^8,9^. That SSPE remains invariably fatal reflects our limited understanding of the neuropathogenic complications of measles.

MeV is a non-segmented single-stranded negative sense RNA virus that is considered a prototypical paramyxovirus^10^. Its genome encodes 6 genes that give rise to 8-9 proteins. The nucleocapsid (N) encapsidates the RNA genome forming RNAse-resistant ribonucleoproteins (RNPs) during viral replication. The phospho-(P) and large (L) proteins form the RNA-dependent RNA polymerase (RdRp) complex that act as a viral transcriptase (P-L) or replicase (N-P-L) at appropriate points in the viral life cycle. The matrix (M) protein facilitates the assembly and budding of the RNP genome from the plasma membrane into virions that contain the fusion (F) and receptor binding proteins (RBP, formerly termed H). All paramyxoviruses require the co-ordinate action of F and RBP to mediate membrane fusion ^11,12^. Some paramyxoviruses like MeV are preferentially cell-associated, can spread cell-to-cell, and efficiently form multi-nucleated giant cell syncytia in appropriate receptor-positive cells^13^.

Primary MeV strains use CD150 and nectin-4 on immune and epithelial cells, respectively^14,15^, neither of which are expressed on neurons or other brain parenchyma cells. This adds to the mystery of how MeV establishes a chronic latent CNS infection that recrudesces many years after recovery from the primary infection. However, characteristic mutations are known to arise in CNS MeV isolates from patients with SSPE or MIBE. Nonsense mutations that result in a non-functional M protein ^16^ and missense mutations that result in a hyperfusogenic F protein ^17,18^ are commonly found. Recombinant MeVs with a functional deletion of the M protein or expressing the hypermutated M protein from an SSPE MeV isolate exhibit enhanced fusogenicity and increased neurovirulence ^19,20^. Similarly, F mutants from neuropathogenic MeV strains also show a hyperfusogenic phenotype in cells that do not express detectable amounts of canonical MeV receptors (CD150 and nectin-4). This *in vitro* hyperfusogenic phenotype is correlated with the ability of neuropathogenic MeV strains to initiate a spreading infection in the CNS *in vivo*, and in human neuronal cell cultures *in vitro* ^21,22 23^. However, syncytia are never observed in the brain or in human neuronal cells. It is unclear how neuropathogenic MeV spreads within the CNS and between neurons without forming syncytia. Proposed mechanisms include the use of a MeV neuronal receptor (although a definitive candidate has not been identified) ^6^, or host factors that could facilitate the putative trans-synaptic spread mediated by the hyperfusogenic F protein ^24^. Nectin-elicited cytoplasmic transfer of MeV^25^ has been proposed as a means to establish the initial transfer of infectious RNPs from epithelial cells to neurons, but not subsequent CNS spread.

Regardless of the underlying mechanism, both MeV-F and - RBP are indispensable for neuronal spread. This suggests that receptor engagement and fusion protein triggering remain essential for MeV spread in the CNS. The convergence of data indicates that the functional hallmark of mutations from SSPE and MIBE strains is the gain of a hyperfusogenic phenotype mediated by MeV-F. Importantly, these hyperfusogenic MeV-F mutants manifest their phenotype most clearly in cells that do not express the canonical MeV receptors (CD150 and nectin-4). For example, a single point mutant from the SSPE Osaka-2 strain (T461I) is able to confer upon wild type (wt) MeV (IC323) the ability to form syncytia and replicate in Vero cells. Similarly, the recombinant IC323-T461I virus can now infect neurons in culture, and cause substantial neuropathology when injected into brains of suckling hamsters ^21^.

The MeV-F protein is functionally constrained^26^ and also well-conserved amongst all clinical isolates^27^. The hyperfusogenic F mutations that neuropathogenic MeV acquires must therefore be highly beneficial to its spread in the CNS. Our ability to understand the fitness landscape of such MeV-F mutations is currently limited by the relatively small number of MeV-F mutations reported to exhibit such a hyperfusogenic SSPE phenotype ^6, 28^. A comprehensive account of the mutational spectrum that can give rise to this hyperfusogenic phenotype will facilitate a better understanding of how these MeV-F mutations confer their fitness advantage.

In this study, we generated a saturation point mutagenesis library of MeV-F in its genomic context. We then designed a fitness screen where only viral genomes bearing MeV-F mutations that mimic the hyperfusogenic SSPE phenotype will have a selective advantage. We not only identified a number of MeV-F mutations similar to ones that have already been reported ^6^, but also numerous novel mutations that span all three structural domains of MeV-F ^29^. Structure-function studies confirm the SSPE phenotype of these mutations and identified a new site on the MeV-F trimer that regulates MeV fusion activity. Finally, we identified a hyperfusogenic mutation in a highly conserved residue in the fusion peptide of MeV-F that was generalizable to Nipah virus, a member of the only other paramyxovirus genus known to harbor viruses that cause chronic latent CNS infections.

## Result

### Saturation mutagenesis screen for MeV-F mutants with hyperfusogenic SSPE phenotypes

#### Rationale

The cardinal phenotype of hyperfusogenic MeV-F mutants from SSPE and MIBE strains is the ability to pair with wild type MeV-RBP and form syncytia in cells that do not express the canonical MeV receptors. In order to better understand the mechanistic features that underlie the hyperfusogenic phenotype of such MeV-F mutants, we designed a screen of a saturating genomic MeV-F mutant library using BSR-T7 cells where this phenotype was the most apparent and rMeV could be rescued with high efficiencies ^26^. As shown in Fig. 1, a rMeV bearing the well characterized F-T461I hyperfusogenic mutant derived from the Osaka-2 SSPE strain^30^ was able to infect, spread and replicate in BSR-T7 cells whereas its isogenic wild type (wt) rMeV (IC323-GFP) counterpart could not. The latter confirms that BSR-T7 cells, a hamster-derived cell line, do not express the canonical MeV receptors as wt rMeV-IC323 replicates well and forms obvious syncytia in cells that express human CD150 or nectin-4^31^. Syncytia and spread (GFP counts) was readily observed with the control rMeV-F T461I hyperfusogenic mutant but not its isogenic wt counterpart (Fig 1A and B). We observed significant differences as early as 2 days post-transfection/rescue. RT-qPCR for genome copy numbers confirmed productive replication of rMeV-F-T461I (Fig 1C), which produced several thousand-fold more viral genomes than wild type rMeV at day 8 post-rescue.

**Figure 1.**
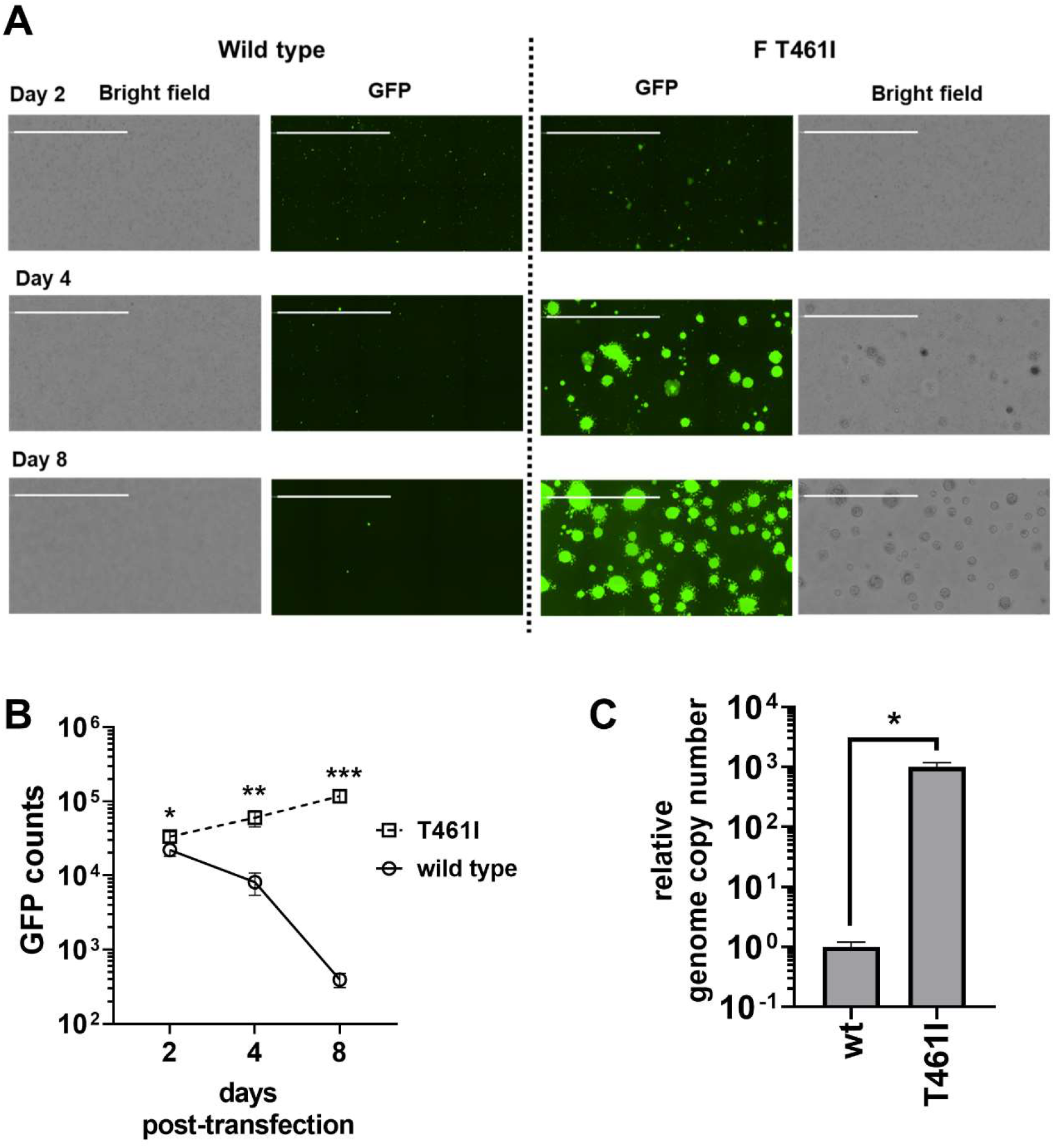
BSR-T7 cells allowed efficient spread of hyperfusogenic (F-T461I mutant) but not wild type (wt) MeV. Recombinant MeV genomes (IC323-GFP) bearing wt F or F-T461I were rescued via reverse genetics in BSR-T7 cells using a transfection-only protocol (see Methods). (A) At 2 days post-transfection (dpt), many GFP+ single cells were seen in both wt and F-T461I rMeV rescue wells. By 8 dpt, most of the GFP+ cells in the wt rMeV rescue wells had disappeared while the GFP+ cells in the rMeV-F-T461I rescue wells showed an increase in both number and size. The latter is indicative of a hyper fusogenic SSPE phenotype. (B) GFP+ cells/syncytia in both the wt and T461I rMeV rescue wells were imaged and quantified on 2, 4 and 8 dpt. Results shown are mean values (+/- standard deviation) from 5 independent experiments. (C) Genome copy numbers in the cytoplasm for both viruses were quantified by RT-qPCR and normalized to the mean value of wt rMeV. Results shown are mean relative genome copy numbers (+/- standard deviation) from 5 independent experiments. Statistically significant differences in the growth and spread of rMeV bearing wt versus F-T461I are indicated (*, p<0.01; **, p< 0.005; ***, p < 0.001, unpaired Student’s t-test). All images shown were captured on the Celigo imaging cytometer (Nexcelom) and GFP+ counts enumerated with the manufacturer’s software. Images are computational composites from an identical number of fields in each rescue well. White scale bar equals 2 millimeters.

#### Library Preparation

Next, we used error-prone PCR to generate a saturating mutagenesis library that covered the entire F gene. We reasoned that only mutants that result in the hyperfusogenic SSPE phenotype exemplified by rMeV-F-T461I would have a fitness advantage when rescued in BSR-T7 cells (Fig. 1C). In order to rescue this MeV-F library in its genomic context and have the requisite coverage (Table S1), we generated four independent mutagenesis libraries of equivalent sizes (402-418 nucleotides) that altogether span the entire MeV-F gene (Fig. 2A). The size limitation of each library (~400 nt) was imposed by the maximum amplicon size we could sequence confidently on Illumina machines (250 nt paired-end reads). The cloning strategy to shuttle each mutagenesis library into the genomic context of rMeV-IC323 is depicted in Fig 2B. Before embarking on making all four libraries, we randomly chose library 3 (MeV-F ORF, nt 840-1241) to optimize our error-prone PCR conditions. Fig 2C summarizes the distribution of mutation rates we observed when we sequenced library 3 generated from low, medium and high error-prone PCR conditions. Sequencing the same region amplified with high fidelity DNA polymerase served as our background control. The mutations were evenly distributed across this library 3 region regardless of mutation rates (Fig S1A to D), which averaged 0.04% for plasmid DNA (background), and 0.12%, 0.18%, and 0.27% for the low, medium, and high mutation settings, respectively (Fig S1E). The mutational spectrum was also relatively unskewed with a transition/transversion (Ts/Tv) ratio close to 1 (0.81). We chose the high error rate condition to move forward with all 4 libraries because we aimed to introduce 1 mutation/400 bp (0.25%), which was the size of each library segment. When we set the threshold detection limit for mutations at 0.08% (double the sequencing error rate (0.04%) of the negative control in Fig S1E) and used the high ‘mutational setting’ for our error-prone PCR conditions, 97.3% of the region covered in library 3 were mutated.

**Figure 2.**
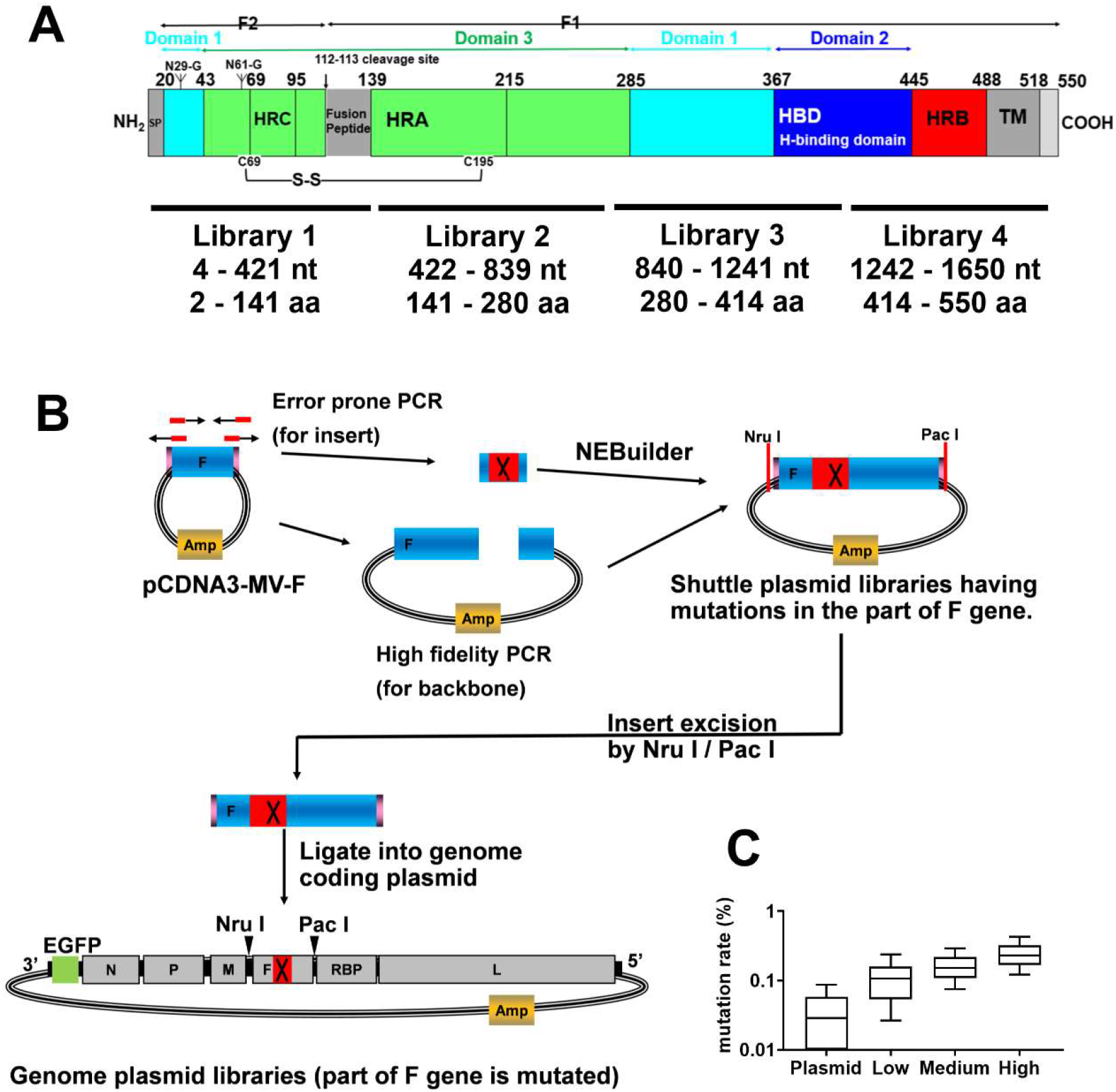
Preparation of genomic MeV-F saturation mutagenesis libraries. (A) Schematic of MeV-F showing the relevant structural and functional domains. Regions targeted by libraries of 1-4 are indicated below the schematic. Nucleotide (nt) and amino acid (aa) positions in the MeV-F open reading frame (ORF) are indicated counting from the initiator methionine. (B) The strategy for using error-prone PCR to construct the MeV-F saturating mutagenesis library in its genomic context is shown. Point mutations in a given library region of F gene were introduced first by error-prone PCR (red X box) and transferred back into the pcDNA3-MV323-F shuttle plasmid via NEB builder. The full F ORF, now containing the mutated F library region, and flanked by NruI/PacI restriction sites, was then transferred to the MeV genome coding plasmid via direct ligation using the same unique restriction sites in the UTRs flanking the MeV-F gene. In this way, four independent measles genome plasmid libraries were generated with saturating F mutations that altogether cover the entire gene. (C) The distribution of mutation rates in pcDNA-MV323-F libraries (library 3) generated under low, medium or high error-prone PCR conditions, or amplified with high fidelity DNA polymerase from the parental ‘plasmid’ is shown. Box and whiskers plot shows the 10/90 percentile (whiskers), 25/75 percentile (box), and median (horizontal bar).

When the rest of libraries (1, 2 and 4) in pCDNA3-MV-F were made with the same high mutation setting, there was also no obvious skewing in the distribution and spectrum of mutations (Fig S2A to C). The mutation rate of libraries 1 (0.33%), 2 (0.27%), and 4 (0.22%) were similar to that of library 3 (0.25%), and all had Ts/Tv ratios close to 1 (Fig S2D). Then, each of the F gene mutated libraries were independently transferred into the genome coding plasmid using the cloning strategy shown in Fig. 2B. Sequencing of the MeV-F libraries in the MeV genome showed that the mutation rate and spectrum were maintained in all libraries with the exception of library 4, where for unknown reasons, the average mutation rate dropped from 0.22% to 0.14% (Table S1). Given that we aimed for one mutation/400 bp and that saturation point mutagenesis requires ~1600 distinct mutations/library, the number of independent genomic clones we generated per library (range: 2.32 – 8.55 x10^4^) ensured that every possible nucleotide substitution at every position was adequately represented. This was confirmed by deep sequencing our genomic libraries where as expected, the coverage was ≥99% for all libraries except library 4 (Table S1).

### Identification of putative hyperfusogenic MeV-F genomic clones in our library screens

We then used our efficient reverse genetics system to rescue wt MeV (IC323-GFP), the four MeV-F mutated libraries, and the hyperfusogenic rMeV-F-T461I (positive control) in BSR-T7 cells as described in Fig. 1^26,32^. The virus producer cells were grown for 8 days with one passage at day 4. The average number of rescue events in each of these screening trials were around 60,000-90,000, which is sufficient to represent the mutational diversity present in each library (Table S1). When wt rMeV-IC323 was rescued and passaged under these conditions, only single cells turned EGFP+, which did not spread. In contrast, libraries 1-4 each gave rise to obvious foci of spreading EGFP+ syncytia by day 8 (Fig. 3A). Each of these large spreading syncytia likely represents a hyperfusogenic mutant similar to that seen with the MeV-F-T461I positive control. These phenotypic observations were confirmed and extended upon analysis of F gene sequences by next generation sequencing NGS at 8 days post rescue. While the genome plasmid libraries contained a relatively even distribution of mutations, we detected a clear selection of fit mutants in all four MeV-F libraries at day 8 post-rescue (Fig. 3B).

**Figure 3.**
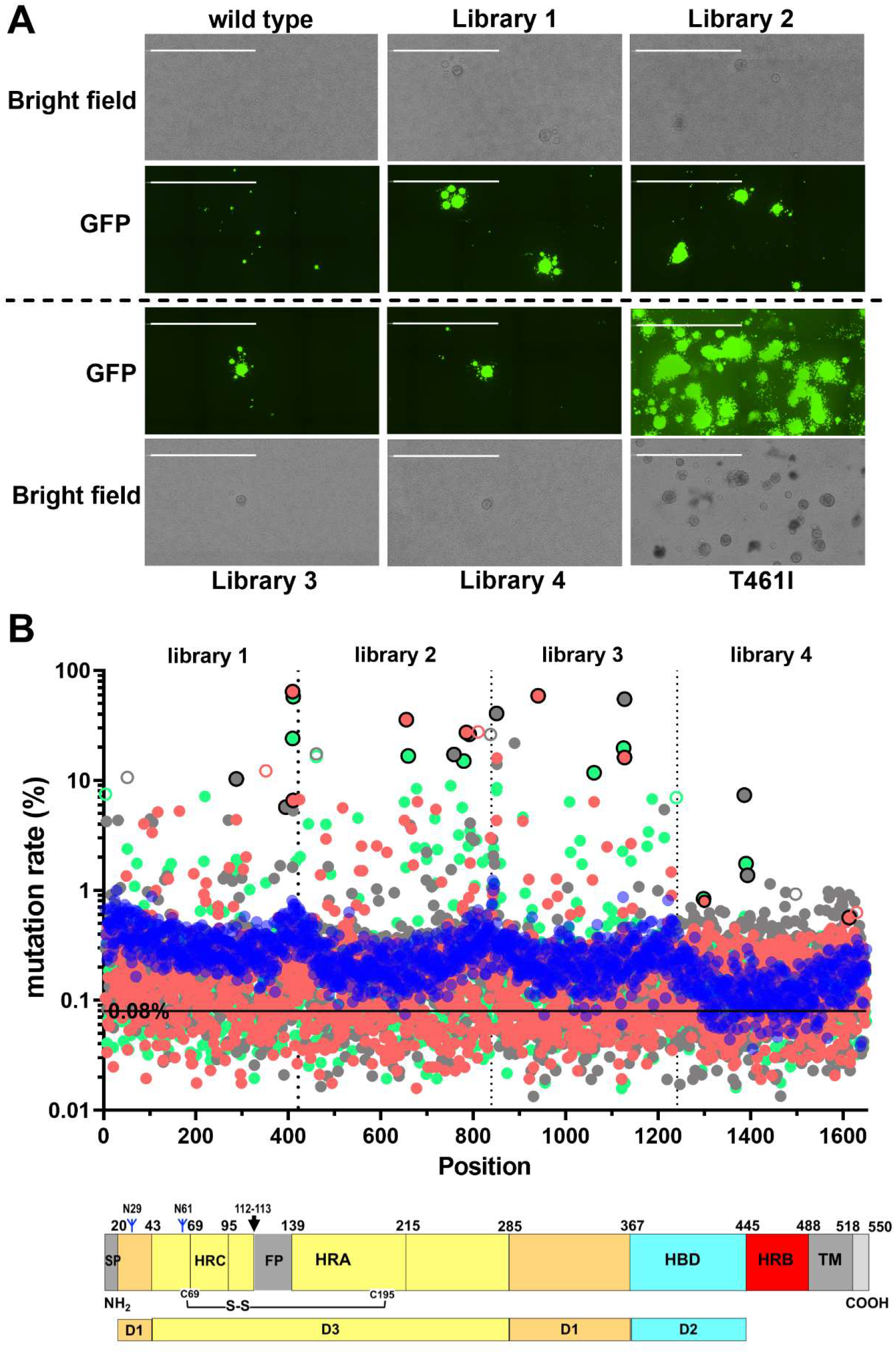
Fitness screen of MeV-F mutagenesis libraries identifies hyperfusogenic F mutants. (A) Representative images from rescue of MeV-F libraries at 8 dpt in BSR-T7 cells show spreading GFP+ syncytia of varying sizes in all libraries except for wt MeV, suggesting that hyperfusogenic mutants are being positively selected. The T461I rescue well showed multiple giant syncytia as expected since only a single clone was transfected. Images were captured by the Celigo Imaging Cytometer (Nexcelom) and is a computational composite of several identical fields of view taken in each well. White scale bar equals 2 millimeters. (B) Mutation rate across the F gene in our saturation mutagenesis genome plasmid libraries before transfection and rescue are indicated by the solid blue circles. The horizontal line at 0.08% represents the threshold we used to define a genuine mutation at any given position. Distribution statistics of the mutants in each genomic library is given in Table S1. Salmon, grey, and green solid circles represents the mutation rate for each library rescued three independent times (replicates 1-3 for library 1-4 = 12X total) based on MeV-F genomic RNA extracted from BSR-T7 cells at 8 days post-rescue. The graph is a concatenation of deep sequencing results of library 1, 2, 3, and 4. The salmon, grey and green circles outlined in black represent the two most predominant mutants in each library and replicate as indicated in Table S2. These mutants were chosen for functional validation in Fig. 4. The open circles correspond to synonymous passenger mutations that were associated with their cognate hyperfusogenic mutants (detailed in Table S2 notes). A scaled schematic of the MeV-F protein is shown below the graph for interpretative convenience.

Each library was independently passaged three times for a total of 12 replicates that made up the screen of the entire MeV-F gene. Some mutants were reproducibly selected in more than one replicate (e.g. L137H, L137F) and/or can account for more than 50% of the library reads at that position (e.g. G376V, T314P, L137F). Table S2 shows the top hit list from these screening experiments. Mutants selected from each library in each replicate were ranked by their percent representation at that position. Despite some stochasticity, it was clear that some aa positions or microdomains were hotspots for mutations that putatively conferred a hyperfusogenic phenotype. This was underscored by the G376V mutant in library 3, which not only dominated the outgrowth of mutants in replicate 1 and 2, but also a ‘fourth’ replicate performed during our preliminary optimization experiments (Table S3). The synonymous mutations that showed up as top hits in Table S2 (e.g. nt 351 mutant in replicate 1 of library 1) were almost always on the same reads of other putative hyperfusogenic mutants, suggesting that these were ‘passenger’ mutations.

Library 4 appeared to be an outlier in that the top hits in all three replicates never exceeded 2% except for N462K in replicate 2 (7.3%). The relatively low mutation rate in the genome library (0.14% vs 0.25-0.35% for libraries 1-3, Table S1) might have reduced the efficiency and spread of mutants that would have otherwise been detected as hyperfusogenic. Remarkably, we were still able to detect and select for previously reported hyperfusogenic mutants such as N462K ^33^ and N465S ^21^ in addition to new mutations like G464W.

### Selected MeV-F mutants have a hyperfusogenic phenotype

We developed a quantitative image based fusion assay (QIFA) to evaluate the large number of potential hyperfusogenic MeV-F mutants identified in our screen. QIFA is premised upon detecting syncytia frequency where syncytia is defined by statistically robust criteria. We first transfected Lifeact-EGFP alone into BSR-T7 cells and obtained a size distribution of single cells in the absence of any syncytia (Fig S3A). We observed that 13 pixcel^2^ was the median size of single cell populations (n=2,605) and GFP objects ≥260 pixcel^2^ (20X median single cell size) was rare in that more than 99% of cells imaged were <260 pixcel^2^. So, we defined a bona fide syncytia as having ≥260 pixcel^2^ and calculated syncytia frequency as a percent (%) of total GFP counts. To help us identify syncytia, we transfected increasing amounts of plasmids expressing MeV-RBP and F-T481I along with a fixed amount Lifeact-EGFP. Our QIFA was highly specific and quantitative, but the QIFA metric (syncytia frequency %) had a dynamic range that plateaued around 8-10% (Fig S3 B and C). The massive syncytia that formed in a finite area (e.g. between 50-100 ng MeV-RBP/F-T461I transfected) limits the numerator. Nonetheless, the assay could be made more or less sensitive by simply increasing or decreasing the hours post-transfection (hpt) before syncytia are quantified (see below).

Next, we chose the best 2 non-synonymous mutants from each individual experiment in Table S2 (Library 1: Q96P, T132S, L137H, L137F; Library 2: P219T, S220I, G253E, L260S, S262N, G264E; Library 3: A284T, H297Y, T314P, L354P, F375L, G376V; Library 4: G433E, S434G, N462K, G464W, N465Y, D538G) plus two mutants (F375V, I393M) from the preliminary screen of library 3 (Table S3). We also included D481G mutant (third best mutant in replicate 2 of library 4) because this corresponds to stalk region in which no previous hyper-fusion mutant was reported. We tested fusion activity of these mutants in comparison with wt F protein which remained mostly as single cells. Several mutants (L137H, L137F, S220I, G253E, S262N, G264E, T314P, L354P, G376V, G464W) showed greatly increased fusion activity (≥ 8%) comparable to the positive control T461I mutant (Fig 4A and 4B) at 30 hpt. Some (Q96P, H297Y) showed moderately increased fusion activity (< 8%). Others (P219T, L260S, A284T, F375V) showed only slightly increased syncytia frequency, which only became statistically significant from wt at 48 hpt (Fig 4C and 4D).

**Figure 4.**
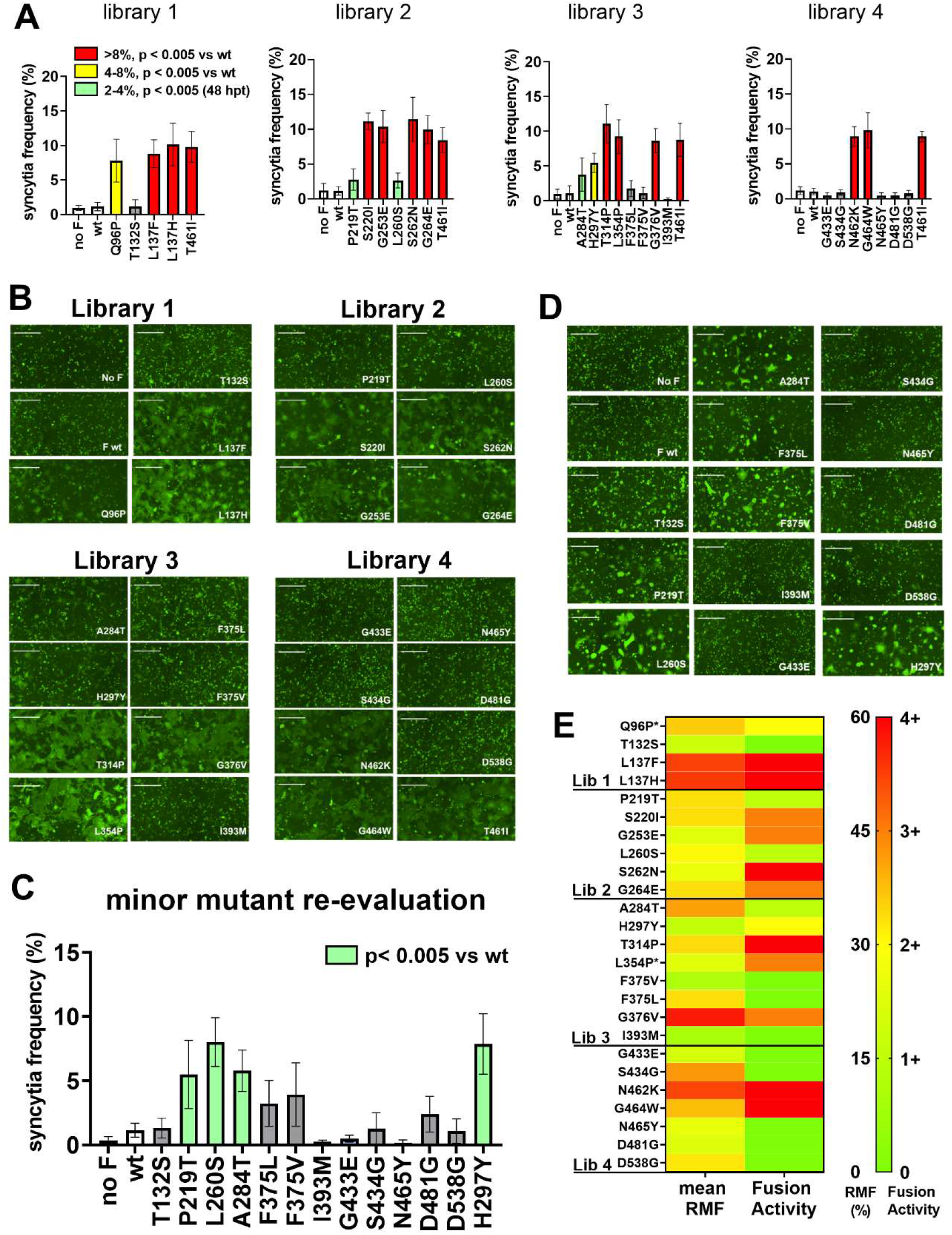
Validation of selected hyperfusogenic MeV-F mutants by a quantitative image based fusion assay (QIFA). We selected the top two mutants from each replicate experiment across all 4 libraries (Table S2) to assess their fusion phenotype (see text for details). We transfected MeV-F/RBP-F and Lifeact-EGFP into BSRT-T7 cells and quantified syncytia formation via our QIFA as described in Fig. S3. (A) Standard QIFA using 200 ng each of MeV RBP and the indicated MeV-F. Images were captured at 30 hours post-transfection (hpt) and analyzed. Data shown are mean (+/- S.D.) syncytia frequency (%) per total GFP counts from five independent experiments. Red, yellow and green bars indicate syncytia formation at the levels indicated in the key. Dunnet’s multiple comparison test was used for the detection of statistically significant differences above wild type MeV-RBP and F. Mutants from each library were assayed with MeV-F-T461I always serving as a positive internal control. Representative images of the summary data in A are shown in (B). For mutants that did not show significant syncytia at 30 hpt, we re-evaluated them at 48 hpt (C). Representative fusion images of C are shown in (D). A heat map summarizes how well the putative hyperfusogenic mutants identified in our library screens correspond to their fusogenic activity (E). Since each library had a different average mutation rate (Table S1) and each replicate was independently rescued and passaged, we first calculated the relative mutation frequency (RMF) for each of the top-ranked mutants (Table S3). RMF = the mutation rate for a given mutation/highest mutation rate for that experiment. In this way, the highest ranked mutant in each library for a given replicate was always 100% and all mutants in that experiment were enriched relative to that highest ranked mutant. The RMF for the indicated mutants from all three replicates can thus be averaged and shown as mean RMF (left column). The fusion activity for the same mutants was categorized into five groups (right column): 0, same as wild typ; 1+, syncytia visible only after 48 hours; 2+, 4-8% at 30 hpt; 3+, ≥8 % at 30 hpt; 4+, significantly different from wt at low expression conditions (25 ng MeV-RBP/F transfected) (Fig. S4). Images shown were generated by the Celigo Imaging Cytometer as described in Fig. 3. White scale bar (B and D) equals 500 micrometers.

To further differentiate between all the hyperfusogenic mutants that gave ≥ 8% syncytia frequency, we repeated our QIFA on these mutants using half the amount of transfected MeV-F/RBP (25 ng each). Under these limiting conditions, 6 of the 11 mutants still showed significant syncytia formation above wt (Fig. S4A and B), as did our positive control T461I mutant. These mutants were located in the C-terminus of the fusion peptide (L137F, L137H), at the protomer interface (S262N), and in the neck region between the head and stalk domains (T314P, N462K, G464W).

Although there was a general correlation between the relative mutational frequency of the dominant mutants identified in the screening experiments and their fusogenicity as measured by our QIFA, not all the mutants identified in the screening experiments were hyperfusogenic (Fig. 4E). Nonetheless, we were able to identify at least 15 novel MeV-F mutations that had a hyperfusogenic SSPE phenotype, effectively doubling the list of hyperfusogenic SSPE-like mutants to date (Table S4).

### A hyperfusogenic mutation in the conserved fusion peptide region is generalizable to the Nipah virus F protein

L137F and L137H mutants repeatedly showed up and were often the predominant mutations in all three replicates of library 1 screens (Table S2). In addition, our quantitative image-based fusion assay (QIFA) revealed L137F and L137H to be as hyperfusogenic as the positive control T461I mutant (Fig. 4A). L137 in MeV-F is located towards the C-terminus of the fusion peptide. The homologous leucine is conserved amongst all major paramyxoviruses (Fig 5A), as demonstrated by our alignment of the fusion peptide region from the indicated viruses. We chose an overrepresentation of F proteins from morbilliviruses and henipaviruses as paramyxoviruses that use protein-based receptors may have differential features for fusion activation^34^. Nonetheless, a phylogenetic tree shows that we chose prototypical viruses that span the diversity within the *Paramyxoviridae* (Fig 5B). We speculated that mutation in this position may also change the fusion activity of other paramyxoviruses, particularly those of henipaviruses, the only other genus of paramyxoviruses known to use protein receptors and also cause chronic latent CNS infections^35^. Thus, we introduced these mutations into NiV-F (L134H and L134F) and evaluated their fusion activity (when co-transfected with NiV-RBP) by our QIFA. L134F markedly enhanced the fusion activity of NiV-F beyond wt whereas L134H did not (Fig 5C and 5D). Interestingly, L137F was also significantly more hyperfusogenic than L137H when fusion was evaluated under limiting conditions (Fig S4A and B) (Tukey’s multiple comparisons test, adjusted p = 0.014).

**Figure 5.**
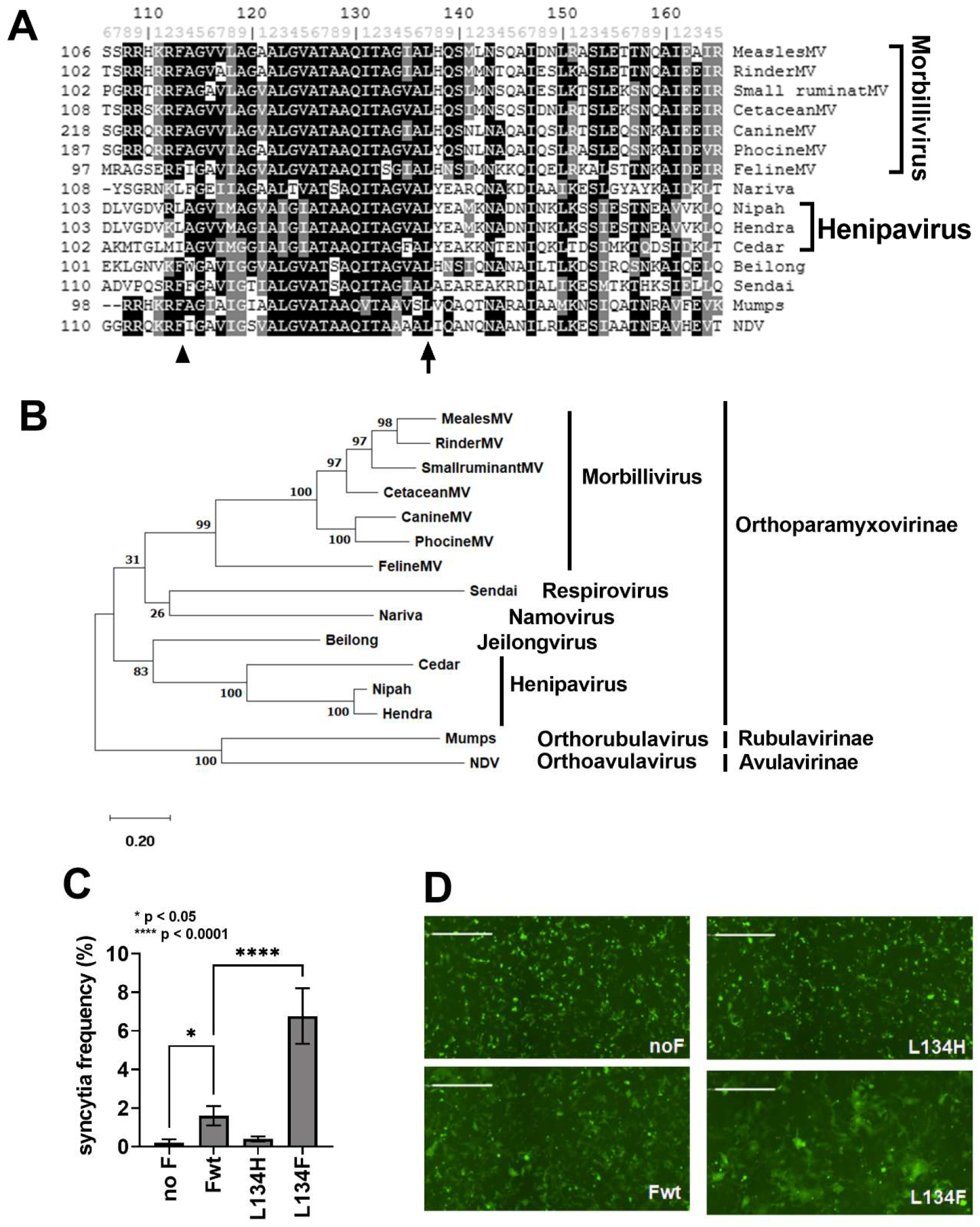
The homologous mutation of MeV-F-L137F in Nipah virus F (NiV-F-L134F) renders NiV-F hyperfusogenic. (A) Amino acid sequence alignment of the fusion peptide region flanking the L137 residue in MeV (arrow) shows that it is highly conserved amongst paramyoviruses. The F_1/F2_ protease cleavage site (arrowhead) is indicated as a point of reference. Prototypic viruses were chosen to represent paramyxoviruses from all three subfamilies and the major genera within each subfamily. (B) A phylogenetic tree of the F protein sequence demonstrates that the selected F proteins span the diversity within *Paramyxoviridae*. Amino acid sequences were aligned by clustalw, and the phylogeny was generated by the maximum likelihood method using MEGA 10 (version 10.1.8). The numbers at the node indicates the fidelity by bootstrap test (1,000 times). The scale indicates substitutions per site. (C) Introduction of the homologous mutation at this position (L134F) made the NiV-F protein hyperfusogenic. Fusion activity In NiV-RBP and F transfected cells was evaluated by our quantitative image based fusion assay (QIFA) as described in Fig. 4. Mean values of 5 independent experiments are shown with error bar indicating standard deviation. Dunnet’s multiple comparison test was used for tests of significance for the indicated comparisons. (D) Representative images of data presented in (B). Images were generated by the Celigo Imaging Cytometer as described in previous figures. White scale bar equals 500 micrometer. Canine morbillivirus (Canine MV); NP_047205, Rinder morbillivirus (Rinder MV); YP_087124, Small ruminant morbillivirus (Small ruminant MV); YP_133826, Phocine morbillivirus (Phocine MV); YP_009177602, Cetacean morbillivirus (Cetacean MV); NP_945028, Feline morbillivirus (Feline MV); YP_009512962, Sendai virus (Sendai); NP_056877, mumps virus (mumps); NP_054711, Nipah virus (Nipah); NP_112026, Hendra virus (Hendra); NP_047111, Cedar virus (Cedar); YP_009094085, Newcastle disease virus (NDV); YP_009513197, Nariva virus (Nariva); YP_006347587, Beilong virus (Beilong); YP_512250.

### Structural mapping of hyperfusogenic MeV-F mutations

Extant hyperfusogenic mutations in MeV-F can be categorized into 3 sites based on how they mapped onto the crystal structure of stabilized trimeric MeV-F ^29^. Site I mutations are in the region surrounding fusion peptide, site II mutants localize to the interface of the protomers while site III mutations cluster in the neck domain between head and stalk. Most of our hyperfusogenic mutants mapped to one of these sites (Fig 6A). Mutants mapping to site I (Q96P, L137F, L137H, F375V, and G376V, Fig 6B) can potentially affect fusion peptide exposure and the biophysical properties of the fusion peptide itself. Site II mutants such as P219T, S220I, G253E, L260S, S262N, and G264E (Fig 6C) can disrupt the inter-protomer interactions that keep F from being pre-maturely triggered. Site III mutants (T314P, L354P, G464W) at the base of the head domain (Fig 6D) can also destabilize the F trimer resulting in lower activation energy for fusion triggering. Interestingly, two of our hyperfusogenic mutants, A284T and H297Y, are located on the beta-sheet connecting the head and neck region (corresponds to Y277 to G301). This region is structurally distinct from sites I - III and no hyperfusogenic SSPE-derived mutants have been mapped to this beta-sheet. We propose classifying this region as site IV, which may represent a novel fusion-regulatory domain encompassing at least the two mutants identified in our study (Fig 6D, A284T and H297Y).

**Figure 6.**
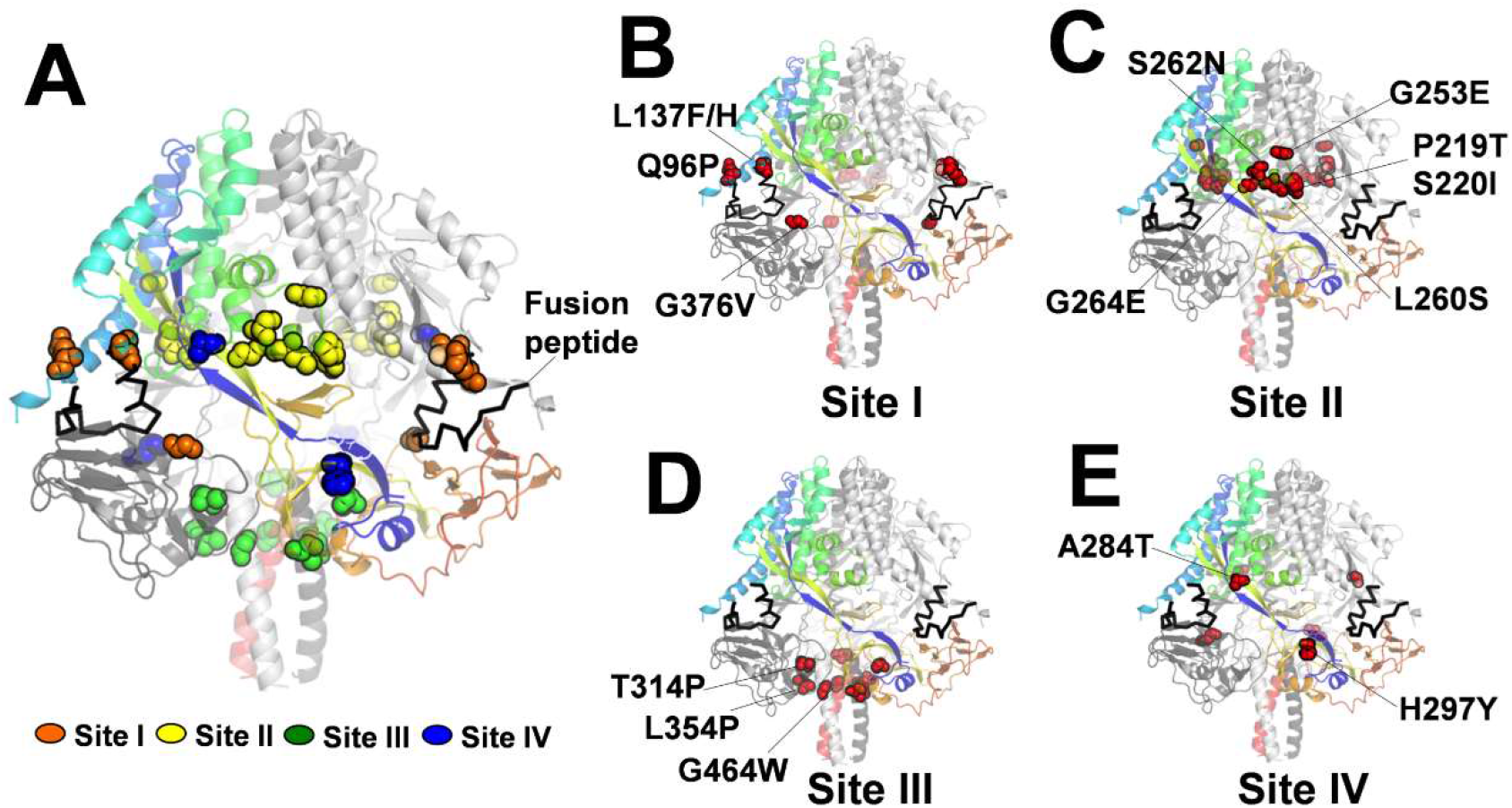
Structural mapping of hyperfusogenic F mutants. (A) Amino acid residues (spheres) whose substitutions confer hyperfusogenicity were mapped onto the trimeric pre-fusion MeV-F structure (PDB:5YXW) using Pymol. A ribbon model is shown where each of the protomers is colored rainbow, dark gray and light gray, respectively. (B-D) The majority of mutants discovered in this study, while novel, mapped to the three sites (I, II and III) that were previously used to classify extant hyperfusogenic mutants by Hashiguchi et al. (E) Two mutants, A284T and H297Y, mapped to structurally distinct beta-sheet that connects the head and neck domains, which we term site IV. In the aggregate model showing all the mutants (A), Site I-IV mutants are represented by orange, yellow, green, and blue spheres, respectively.

### Recombinant measles virus expressing selected hyperfusogenic F mutants recapitulate the SSPE phenotype

To confirm that the hyperfusogenic MeV-F mutants identified in our screen was necessary and sufficient to confer a SSPE phenotype, we chose 4 mutants (L137F, S262N, H297Y, and G464W) to rescue as isogenic rMeV. We chose the best mutant from sites I-IV as evaluated by our QIFA (Fig 4A and Fig S4A). For site III, we chose G464W instead of N462K because N462K has already been reported^33^. rMeV-IC323^EGFP^ with the L137F, S262N, or G464W F mutations formed huge syncytia (Fig 7A) and replicated several hundred-fold better than wt even at day 6 post-rescue (Fig 7B). rMeV with the F-H297Y mutant showed a less dramatic increase in the syncytia size at day 6, but nonetheless showed significant increase in genome copy numbers by 8 days post-rescue compared to wt (Fig 7C). To substantiate the biological relevance of our structural mapping efforts, we used a well-characterized fusion inhibitory peptide (FIP) that not only blocks MeV fusion ^36^, but does so specifically by interacting directly with residues in site III such as G464 ^29^. We found that the G464W mutant was resistant to FIP but not the other non-site III mutants (Fig 7D and 7E). To ensure that these results were not due to differences in surface expression and/or F protein cleavage, we performed cell surface biotinylation experiments. Briefly, surface expressing F proteins were pull downed by streptavidin-beads after cell surface protein biotinylation, and the immunoprecipitated cell surface MeV-F detected by Western blot using a MeV-F specific antibody (Fig 7F). Densitometry showed that surface F protein expression was highest in wt F and the T461I mutant, but all the hyperfusogenic mutants (L137F, S262N, and G464W, and H297Y) were expressed at ~50% of wt levels mutants (Fig 7G). Cleavage efficiency was also evaluated by the analyzing the ratio of F1/F0 (Fig 7H), which showed no significant differences in cleavage efficiency between wt and any of the hyperfusogenic mutants. Altogether, these results show that the hyperfusogenic phenotype was not due to overexpression or more efficient cleavage of the indicated F mutants.

**Figure 7.**
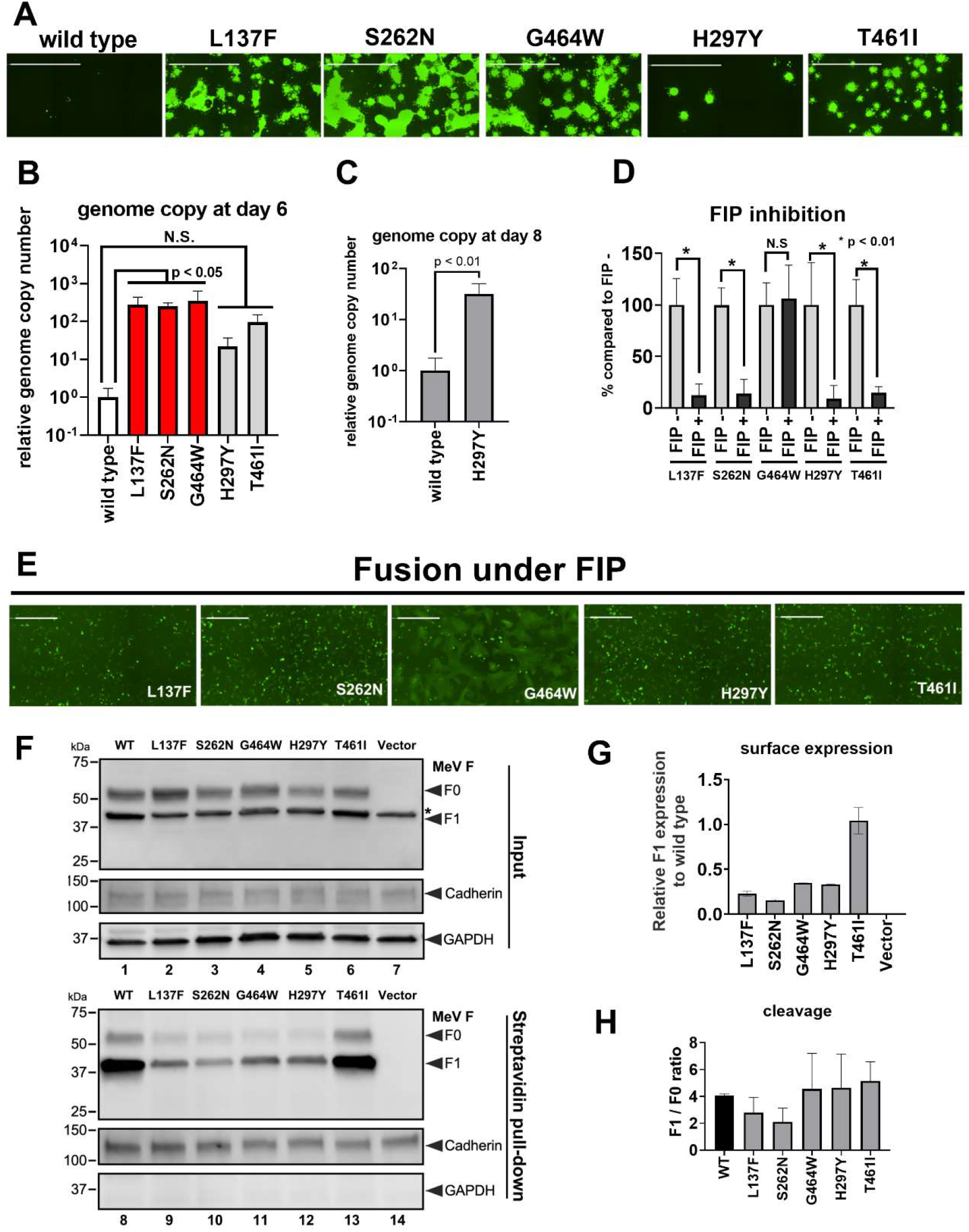
Recombinant MeV expressing the hyperfusogenic F mutants from each of the four sites exhibit the expected phenotype. Recombinant MeV with wt F, T461I, and representative F mutations (L137F, S262N, G464W, H297Y) from sites I-IV (Fig. 6) were rescued in Bsr-T7 cells. Virus growth was monitored via fluorescence microscopy (A). At day 6 post rescue (with one passage at day 3), cytosolic RNA was collected and genome copy number was quantified by RT-qPCR (B). Genome copy numbers for all the mutants were normalized to that for wt MeV, which was set to 1 (B and C). Growth of H297Y mutants was further evaluated for 8 days incubation period (with one passage at day 4) (C). Data shown are mean relative genome copy numbers (+/- S.D) from 5 independent experiments (B and C). Statistically significant differences from wt were determined by Dunnet’s multiple comparison test (B) and t-test (C). Syncytia images were generated by the imaging cytometer as in previous figures. White scale bar equals 2 milimeters. MeV-specific fusion-inhibitory peptide (FIP) inhibited the fusion activity of all the hyperfusogenic mutants except the G464W mutant, as evaluated by our QIFA (D). Data shown are mean (+/- S.D.) from 5 independent experiments. Representative images of the fusion inhibitory data are shown in (E). P<0.05 or <0.01 for the indicated comparisons (Student’s t-test). Images generated on the image cytometer as before. White scale bar equals 500 micrometers. (F and G) Cell surface biotinylation experiments show that the L137F, S262N, G464W, and H297Y hyperfusogenic mutants were expressed at lower levels than wt F or F-T461I. At 48 hpt, biotinylated cell surface proteins on MeV-F transfected 293T cells were pulled down by streptavidin-beads and western-blotted with a MeV-F specific antibody. Cadherin and GAPDH served as respective cell surface and cytosolic protein controls. The upper and lower blots in (F) show the input and surface protein (streptavidin pull-down) fraction, respectively. The full-length F0 and cleaved F1 products are marked (arrowhead). * indicates the non-specific band from the cytosol (upper blot) which disappeared in the cell surface pull-down fraction (lower blot). (G) shows the relative F1 surface expression levels of the various F proteins normalized to wt F set at 1 (based on densitometric measurements). (H) shows the F1 / F0 ratio of cell surface F, which indicates cleavage efficiency. Data shown are the average and range of two independent experiments.

A cardinal feature of SSPE MeV strains is the ability to infect and spread in primary human neurons, which do not express the canonical receptors for primary strains of MeV (CD150 and nectin-4). Wild-type MeV can infect neurons, albeit inefficiently, but neurovirulent SSPE/MIBE MeVstrains can infect and spread in cultured primary human neurons. To mimic primary human neuronal infection, we used human iPSC-NPC-derived neurogenin 2(NGN2)-induced glutamatergic neurons that are a well-characterized model of excitatory forebrain neurons^37^. Upon wt rMeV-IC323^EGFP^ infection, we detected a few GFP+ neurons at 2 days post-infection (dpi). GFP appeared distributed in the cell bodies as well as the dendrities and axons that outlined the neurons distinctly (Fig. 8A). On rare occasions, a small cluster of GFP+ neurons could be found connected by GFP+ neurites, which suggest a slow cell-to-cell spread of MeV (Fig. 8A, middle and bottom panels). This is reminiscent of what was reported by Sato et al. in NT2-derived neurons^22^. In contrast, inoculation of the same neuronal cultures with the rMeV-F-L137F mutant resulted in much larger clusters of GFP+ neurons that sometimes appeared to coalesce (Fig. 8B, top panel). The latter show that novel hyperfusogenic F mutants identified in our screen can recapitulate the SSPE phenotype involving efficient spread in primary human neurons.

**Figure 8.**
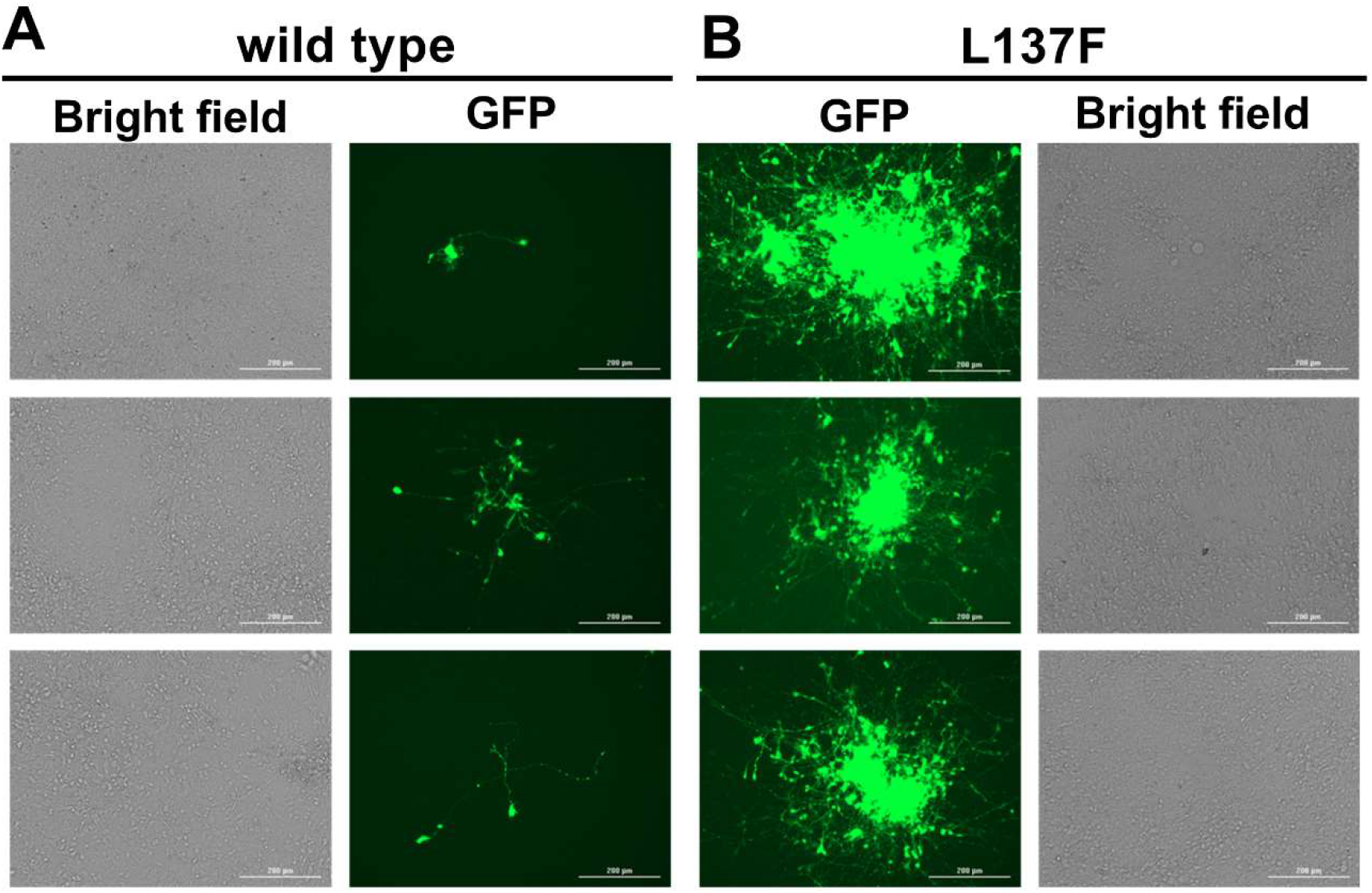
Recombinant MeV with hyperfusogenic F-L137F mutant infects and spreads efficiently in primary human neurons. Homogenous iPSC-derived NGN2-induced glutamatergic neurons, seeded and differentiated in 24-well plates, were infected with 5000 PFU of wild type MeV (A) or MeV-F-L137F (B) per well. (A) At 2 dpi, rare GFP+ neurons could be seen in wild type MeV infected wells. GFP was localized to the cell body as well as the axon and dendrites. Occasionally, small clusters of several neurons could be seen (middle panel). (B) In contrast, MeV-F-L137F infected wells showed numerous GFP positive neurons in large clusters at 2 dpi. 3 representative images from 3 independent infections are shown. Images were taken on the Cytation 3. White scale bar equals 200 micrometers.

## Discussion

Our saturating mutagenesis screen allowed us to interrogate the fitness landscape that allowed for the selection of hyperfusogenic MeV-F mutants associated with SSPE/MIBE phenotypes, which is the ability to pair with wt MeV-RBP and form syncytia in cells lacking the canonical MeV receptors. We identified at least 15 such hyperfusogenic mutants, effectively doubling the list of known hyperfusogenic SSPE/MIBE mutants curated in the past several decades (Table S4). These mutants are a community resource which can be used to shed further light on fusion regulatory mechanism and aid in the search for the putative neuronal receptor for MeV.

The antigenic variability of MeV is highly constrained^38^. All vaccine strains are derived from genotype A, yet sera from vaccinees can neutralize the broad range of circulating genotypes. The monoserotypic nature of MeV implies the lack of antigenic drift and functional constraints on its surface envelope glycoproteins (F and RBP). The latter is highlighted by our previous study showing that MeV F and RBP are completely intolerant to 15 bp (5 aa) insertions during a whole genome transposon mutagenesis screen ^26^. However, the unit of selection in neurovirulent MeV strains appears to be single nucleotide mutations in MeV-F that confer the ability for neuronal spread ^6,21^. This adaptation of MeV to spread within the central nervous system (CNS) is correlated with *in vitro* syncytia formation and cell-to-cell spread in the absence of canonical receptors. Therefore, we prepared saturation point mutagenesis libraries of MeV-F in its genomic context and performed a reverse genetics rescue screen in a hamster derived cell line (BSR-T7), where hyperfusogenic SSPE-like F mutants can spread and replicate, but not wt MeV (Fig 1). The use of the well characterized F-T461I SSPE mutant helped us in choosing the optimal cell line for the screen (Fig. 1) and served as a positive control for each replicate (Fig. 3) and functional validation (Fig. 4).

For saturation coverage, we prepared four separate libraries of ~400 nt in size that altogether span the entire F gene (Fig. 2). There are biological and technical reasons for this strategy. To ascertain single mutations that can act independently in a given library, the library size is limited by the maximal Illumina read depth (2x 250 bp paired end reads), which is effectively <450 bp if one excludes the primer binding sites. To maximize the quality of our library and sensitivity of our screen, we also wanted the biggest difference between the average mutation rate of our libraries and the intrinsic error rate of Illumina sequencing, which we confirmed to be 0.04% in our hands (Fig. 2C, Table S1E). Illumina’s reported error rate is between Q35 – Q40 (0.035% - 0.001%). If we had targeted the entire 1.6 kb F gene with an average of one point mutation, the average mutation rate would be 0.063%, which will be statistically indistinguishable from intrinsic error rate of Illumina sequencing. However, introducing a single point mutation in ~ 400 bp will result in an average mutation rate of 0.25%, which is comfortably 8-fold above the background error rate. A similar strategy was also adopted by a previous influenza mutagenesis screening study ^39,40^ where the authors separated the influenza genome into 52 parts comprising of 250 bp segments.

Although some synonymous mutants appeared in our top hit list (Table S2), most were on the same read of other non-synonymous mutations that were later confirmed to be hyperfuosgenic. This suggests that these synonymous mutations are passenger mutations associated with bona fide hyperfusogenic mutations. Separation of the F gene into four independent libraries thus enabled us to designate these synonymous or neutral mutations as such.

The veracity of our functional screen for hyperfuosgenic MeV-F in its genomic context is underscored by our ability to recover most of the extant hyperfusogenic F mutants that exhibit a SSPE phenotype as long as that mutant was present in our original library. For hyperfuosgenic mutants that we did not recover, close inspection of our data (Table S4) revealed that they were either not present in the original genomic mutant library (e.g. L454W), or present but outcompeted by a distinct mutant at the same amino acid residue (e.g. S262N>>S262R, L354P>>L354M, T461A>>T461I, G464W>>G464E), or known to be hypefusogenic only when present together (G168R, E170G, AS440P, R520C and L550P). This lends greater support to the 16 hyperfusogenic mutants we validated from our screening experiments (lib1: Q96P, L137H, L137F; lib2: P219T, S220I, G253E, L260S, S262N, G264E; lib3: A284T, H297Y, T314P, L354P, G376V; lib4: N462K, G464W). Except for the previously characterized N462K mutant, others were all novel mutants.

L137 mutants are particularly interesting. L137 is located in the C-terminus of fusion peptide and is highly conserved among all paramyxoviruses. The homologous L134F mutation in the NiV-F protein also made NiV-F hyperfusogenic in the presence of NiV-RBP (Fig 5C and 5D). How far this extends to other paramyxoviruses is a subject for future studies.

In addition to validating their phenotype in our fusion assays, we rescued four mutants (one from each library) as independent rMeV clones and showed that they exhibited the expected hyperfusogenic SSPE phenotype (Fig. 7). Altogether, these data indicate our saturation mutagenesis screening system is robust and efficiently identifies mutants dictated by the screening conditions. This system can also be applied to genes of interest in other paramyxoviruses to find interferon vulnerable mutations for better vaccine development as was done for influenza ^39^.

A majority of the previously reported hyperfusogenic mutations (Table S4) are located on site III (part of library 3 and library 4 in our study) in the prefusion structure of trimeric MeV-F (Fig.6, and Table S4), which might have overemphasized the contribution of this site (base of the head domain) to the neurovirulent hyperfusogenic phenotype. However, our screening found many novel and strong phenotypic mutations in site 1 and site 2, suggesting that mutations in all sites found across all three structural domains of F can contribute to the hyperfusogenic phenotype. In addition, we found new mutants – A284T and H297Y— which are located in the beta-sheet (Y277-G301) that connect the head and neck domains (Fig 6D). These two mutants in this structurally distinct site IV do not appear to have as strong a phenotype but are nonetheless real as rMeV-F-H297Y can grow and replicate in BSR-T7 cells.

As expected from crystal structure, G464W may resist FIP inhibition because G464 directly interacts with FIP in the complex crystal structure ^29^. Our data showed that G464W was indeed resistant to FIP inhibition (Fig 7E) just as G464E was also reported as a FIP-resistant hyper fusiogenic mutation ^41^. This suggest that our screen can be judiciously applied to study the barriers to drug or antibody resistance.

Our reverse genetics rescue screen based on a particular phenotype is made possible by the transformative improvements in the efficiency of paramyxovirus rescue. This can now be applied to any gene in any paramyxoviruses for which rescue efficiency is high enough.

## Method

### Cell lines

293T cells (ATCC),Vero cells (CCL-81; ATCC), and BSR T7 cells ^42^ were propagated in Dulbecco’s modified Eagle’s medium (Invitrogen) supplemented with 10% fetal bovine serum (Atlanta Biologicals) at 37°C. Vero cells constitutively expressing human SLAM (Vero-hSLAM cells) were gifted from Dr. Yusuke Yanagi ^43^.

### Plasmids and viruses

We modified the genome coding plasmid for MeV (p(+) MV323-AcGFP) ^44^ and generated pEMC-MV323-EGFP; AcGFP gene was deleted and GFP gene were inserted at the head of genome, and hammer head ribozyme was introduced to increase rescue efficiency. This clinical isolate is well characterized and based on the clinical isolate of H4 strain ^45^. Helper plasmid for virus (N, P, and L protein expression plasmid) rescue were gifted from Dr. Richard Plemper ^32^. The plasmid coding full length viral genome were maintained in Stbl2 E. coli (Thermo Fisher Scientific) with growth at 30°C.

DNA fragment of measles RBP and F gene were amplified from p(+) MV323-AcGFP and cloned into pCDNA3 or pCAGGS vector, creating pCDNA3-MV-F, pCAGGS-MV-RBP, and pCAGGS-MV-F. We introduced mutation to the plasmid or modified the plasmid with the site directed mutagenesis by overlapping PCR, followed by in-fusion (Clontech) reaction to fuse the mutated PCR fragment with backbone cut by restriction enzyme. Primer set used for making each mutant are listed in Table S5.

### Error prone PCR based mutagenesis

Point mutations in F gene were introduced by Genemorph II random mutagenesis kit (Agilent) which uses error-prone PCR to introduce mutations. We divided F gene into 4 part with 402 - 418 bps size; library 1 (418 bp), library 2 (418 bp), library 3 (402 bp), and library 4 (409 bp) as written in Fig 2A. Mutation rate can be adjusted by adjusting the amount of initial template. We used DNA fragments from pCDNA3-MV-F cut by Nru I-HF (NEB) and Pac I (NEB) restriction enzyme as an error-prone PCR template. Amplified PCR fragments were fused with backbone prepared by high-fidelity PCR of Cloneamp (Clonetech) using NEBuilder (NEB), then transformed into stellar competent cells (Clonetech). We scaled up so that each library generate > 20,000 colonies. Bacterial colonies were collected 24 hours after plating and then amplified in Terrific Broth medium (Research Products International) for several hours. DNA were extracted by Plasmid DNA Maxi Prep Kits (Thermo Fisher Scientific).

Next, the insert for genome plasmid was made by treating pCDNA3-MV323-F mutated libraries with Nru I and Pac I. The insert and genome backbone (pEMC-MV323-EGFP cut by Nru I-HF and Pac I) were ligated with T4 DNA ligase (NEB), followed by transformation into electrocompetent cells with ElectroMAX stbl4 (Thermo Fisher Scientific), securing > 20,000 colonies in each library. Bacterial colonies were collected 24 hours after plating and then amplified in Terrific Broth medium for several hours. DNA (F gene mutated genome coding plasmid) were extracted by Plasmid DNA Maxi Prep Kits.

### Recovery of recombinant measles virus from cDNA

For the recovery of recombinant MeV, 4 × 10^5^ BSR-T7 cells were seeded in 6-well plates. The next day, the indicated amounts (detailed below) of antigenomic construct, helper plasmids (-N, -P and -L), T7 construct, and Lipofectamine LTX / PLUS reagent (Invitrogen) were combined in 200 uL Opti-MEM (Invitrogen). After incubation at room temperature for 30 minutes, the DNA - lipofectamine mixture was added dropwise onto cells. Transfected cells were incubated at 37°C with medium replacement every day. The amount of measles plasmids used for rescue follows our previous study ^32^: 5 ug antigenomic construct, 1.2 ug MeV-N, 1.2 ug MeV-P, 0.4 ug MeV-L, 3 ug of a plasmid encoding a codon-optimized T7 polymerase, 5.8 uL PLUS reagent, and 9.3 uL Lipofectamine LTX.

### Growth comparison between wild type and F T461I mutant

Plasmid coding wild type measles or F T461I were transfected to one well (6 well plate) of BSR-T7 cells, then cells were grown with medium replacement every day. At day 4, BSR-T7 cells were treated with trypsin, and 1 / 6 cells were passed into one well in new 6 well plate, then grown for additional 4 days. The number of GFP positive cells was counted at day 2, day 4, day 8 by Celigo Imaging Cytometer (Nexcelom). Supernatant was collected for titration at day 4 and day 8. We modified waiting time down to 6 days (cells were passaged at day 3 once) when comparing the growth of L137F, S262N, and G464W, because syncytia became quite huge and detached from dishes for 8 days waiting time.

### Tittering viral supernatants by plaque assay

Rescued MeV were grown in Vero-hSLAM cells, then supernatants were collected. For tittering virus, monolayer of Vero-hSLAM cells in 12 well was infected by 500 ul of serially diluted samples for 1 hour, followed by medium replacement with methylcellulose containing DMEM. 4 days later, the number of GFP positive plaque was counted under fluoro-microscope to decide titer.

### Passaging virus in the screening experiment

MeV genome coding plasmid with mutations in F protein (genome libraries) were transfected to BSR-T7 cells with helper plasmid and T7 expressing plasmid (as indicated at ‘Recovery of recombinant MeV from cDNA’). Rescued viruses were grown in BSR-T7 cells for 4 days with medium replacement every day. At day4, BSR-T7 cells in one well of 6-well plate were treated by trypsin and passed into 6 wells of 6-well plate. Then BSR-T7 cells with MeV were grown for additional 4 days (total 8 days). Cytosolic RNA was extracted, reverse transcribed, and amplified, then sequenced by next-generation sequencer to evaluate variation of F gene sequence after growth in BSR-T7 cells.

### RT-PCR and Illumina sequencing

The cytosolic RNA was extracted by phenol-chloroform method with Trizol (Thermo Fisher Scientific). The entire F gene sequence was amplified by RT-PCR reaction using Superscript III (Thermo Fisher Scientific) with a primer set to detect F gene, creating 1.7 K bp fragment. The RT-PCR fragment were used as a template for nested PCR by Cloneamp to amplify the sequence of mutated part (402 – 418 bp, excluding primer sequence). The information of primer is available in Table S5.

DNA library preparations, sequencing reactions, and initial bioinformatics analysis were conducted at GENEWIZ, Inc. (South Plainfield, NJ, USA). DNA Library Preparation, clustering, and sequencing reagents were used throughout the process using NEBNext Ultra DNA Library Prep kit following the manufacturer’s recommendations (Illumina, San Diego, CA, USA). End repaired adapters were ligated after adenylation of the 3’ends. The pooled DNA libraries were loaded on the Illumina instrument according to manufacturer’s instructions. The samples were sequenced by MiSeq on a 2x 250 paired-end (PE) configuration. Base calling was conducted by the Illumina Control Software (HCS) on the Illumina instrument.

### Data analysis and selection of the top list

Paired-end Fastq files were merged by BBtools to make single read. We trimmed low quality nucleotides in edge of sequence to by PRINSEQ because 5’ and 3’end of sequence tend to show low quality value ^46^. We trimmed nucleotides until PRINSEQ found the first nucleotide of quality value (QV) => 33, then we selected the read with the average QV >= 33.

WIG file (include nucleotide counts at each position) were generated from processed SAM and BAM files with IGV tools. Mutation rate were calculated from nucleotides counts at each position using reference sequence file.

### Genome quantification by qPCR

Extracted RNA was reverse transcribed by genome specific primer (written in Table S5) with Tetro Reverse Transcriptase (Bioline), then the number of genome was quantified by SensiFAST™ SYBR^®^ & Fluorescein Kit (Bioline) and CFX96 Touch Real-Time PCR Detection System (Biorad).

### Image based fusion assay

BSR-T7 cells were seeded at a density of 40,000 cells/well onto 48-well dish at 24 hours before transfection. Then, cells were transfected with 200 ug of pCAGSS-MV-RBP, 200 ug of pCAGGS-MV-F or F-mutants, and 50 ug of pEGFP C1 Lifeact-EGFP (Addgene, plasmid #58470) with 1.25ul of Lipofectamine 2000 (Invitrogen). At 30 hours (for the evaluation of major mutants) or 48 hours (for the evaluation of minor mutants) post transfection, GFP+ cells and syncytia were captured via an imaging cytometer (Celigo™, Nexcelom) using the green channel (Ex/Em: 483/536 nm). Images were exported at the resolution of 5 um / pixel. The GFP-positive foci (single cell or syncytia by fusion) were analyzed by ImageJ (developed by NIH), creating the profile of individual GFP-positive foci with size information.

For quantification of syncytia formation that reflects a hyperfusogenic phenotype, we first measured the size distribution of single cells. Over 5 replicates of cells transfected with MV-RBP and LifeActGFP only (8:1), which should not result in any syncytia, the median size of putative single cells was 13 pixels^2^ (n= 2,605 objects). We then counted all GFP-positive foci ≥ 7 pixel^2^ (half of the median size of single cells). Then we calculated the frequency of syncytia which is defined as the GFP foci of ≥ 260 pixel^2^ (20 times of average median size of single cells) / total GFP counts of ≥7 pixcel^2^.

For Nipah virus fusion assay, we used 100 ng of pCAGGS-Nipah-RBP-HA and 100 ng of pCAGGS-Nipah-F-AU1 (both from Malaysia strain) and waited 30 hours post transfection.

For NDV fusion assay, we used 100 ng of pCAGSS-NDV-RBP, 100 ng of pCAGGS-NDV-F (cloned from lasota strain) and waited 48 hours post transfection.

### Fusion inhibitory peptide (FIP)

FIP (Z-D-Phe-Phe-Gly-OH) was purchased from BACHEM (#4015768). FIP was dissolved into 10mM solution by DMSO. FIP solution was added to the medium at the final concentration of 2% (200 uM) at 3 hours post transfection. DMSO was used as a negative control of FIP.

### Cell surface expression analysis of F protein

12 ug pCAGGS-IC323-F or F mutants (L137F, S262N, G464W, H297Y, and T461I) were transfected to 293T cells (8x 10^6 cells in 10 cm dish) with X-tremeGENE™ HP DNA Transfection Reagent (Roche). 48 hours after transfection, cell surface proteins were labeled with biotin, then pulled down by streptavidin beads by using cell surface biotinylation and isolation kit (ThermoFisher, #A44390). Collected surface expressed proteins were run on 4 - 15% poly polyacrylamide gel (Bio-rad. #4561086) and transferred onto PVDF membrane (FisherScientific, #45-004-113), followed by primary antibody reaction and secondary antibody reaction. For the detection of measles virus F protein, rabbit polyclonal antibody (Abcom, ab203023) was used as the primary antibody. For the detection of internal control, pan-cadherin antibody (Cell signaling, #4068) and GAPDH Rabbit monoclonal Antibody (Cell signaling technology, #2118) were chosen as cell surface and cytosolic protein, respectively. Alexa Fluor 647-conjugated anti-rabbit antibody (Invitrogen, #A-21245) were used as secondary antibody appropriately. Image capturing and signal intensity analysis were done by Chemidoc™ MP (Biorad).

### Neuron infection experiment

On day −1 NPCs were dissociated with Accutase Cell Detachment Solution for 5min at 37°C, counted and seeded at a density of at 6×10^4^ cells/well on Matrigel coated 24-well plates in hNPC media (DMEM/F12 (Life Technologies #10565), 1x N-2 (Life Technologies #17502-048), 1x B-27-RA, 20 ng/mL FGF2 (Life Technologies)) on Matrigel (Corning, #354230). On day 0, cells were transduced with rtTA and *NGN2* lentiviruses as well as desired shRNA viruses in NPC media containing 10 μM Thiazovivin and spinfected (centrifuged for 1 hour at 1000g). On day 1, media was replaced and dox was added with 1ug/mL working concentration. On day 2, transduced hNPCs were treated with the corresponding antibiotic to the lentivirus (1 μg/mL puromycin for *NGN2*-Puro). On day 4, medium was switched to Brainphys neuron medium (Brainphys (STEMCELL, # 05790), 1% N-2, 2% B-27-RA, 1 μg/mL Natural Mouse Laminin (Thermofisher, # 23017015), 20 ng/mL BDNF (R&D, #248), 20 ng/mL GDNF (R&D, #212), 250 μg/mL Dibutyryl cyclic-AMP (Sigma, #D0627), 200 μM L-ascorbic acid (Sigma, # A4403) plus 1 μg/mL dox. From day 6 to day 15, 200nM cytosine arabinoside (Ara-C) was added to Brainphys neuron medium to prevent extended proliferation, medium was changed half every second day. For maturation, medium was replaced half with Brainphys neuron medium every second day until infection on day 21.

5 x 103 PFU /well (tittered in Vero-hSLAM cells) of wild type MeV and MeV-F L137F were added to infect the neurons for one hour, then virus reagent was removed and replaced by fresh medium. Cells were incubated for 2 days, then fixed by 2%-PFA at 2 days post infection. Infected cells were visualized by Cytation 3 (Biotek).

## Data and materials availability

The raw next generation sequencing results are uploaded at NCBI GEO: GSE14767.

## Ethics declaration

All hiPSC research was conducted under the oversight of the Institutional Review Board (IRB) and Embryonic Stem Cell Research Overview (ESCRO) committees at the Icahn School of Medicine at Mount Sinai (ISSMS). Informed consent was obtained from all skin cell donors as part of a study directed by Judith Rapoport MD at the National Institute of Mental Health (NIMH).

## Acknowledgement

This work was supported by Japan Agenct for Medical Research and Development (AMED) Grant 20wm0325002h (to T.H.), JSPS KAKENHI Grant Numbers 20H03497 (to T.H) and Joint Usage/Research Center program of Institute for Frontier Life and Medical Sciences, Kyoto University.”

## Authors contributions

S. I. and B. L. conceived this study. S.I. conducted library preparation, screening experiment, fusion assay, and virus growth analysis. T. H. did the structural discussion of measles F protein. C. H. conducted the surface expression analysis. K. R., and K. B. worked on human iPS cells derived neuron experiment. M. T. provided measles genome coding plasmid in this study. B. L. supervised this study. S.I. and B.L wrote the manuscript.

