## Supplement information for "Fitness selection of hyperfusogenic measles virus F proteins associated with neuropathogenic phenotypes"

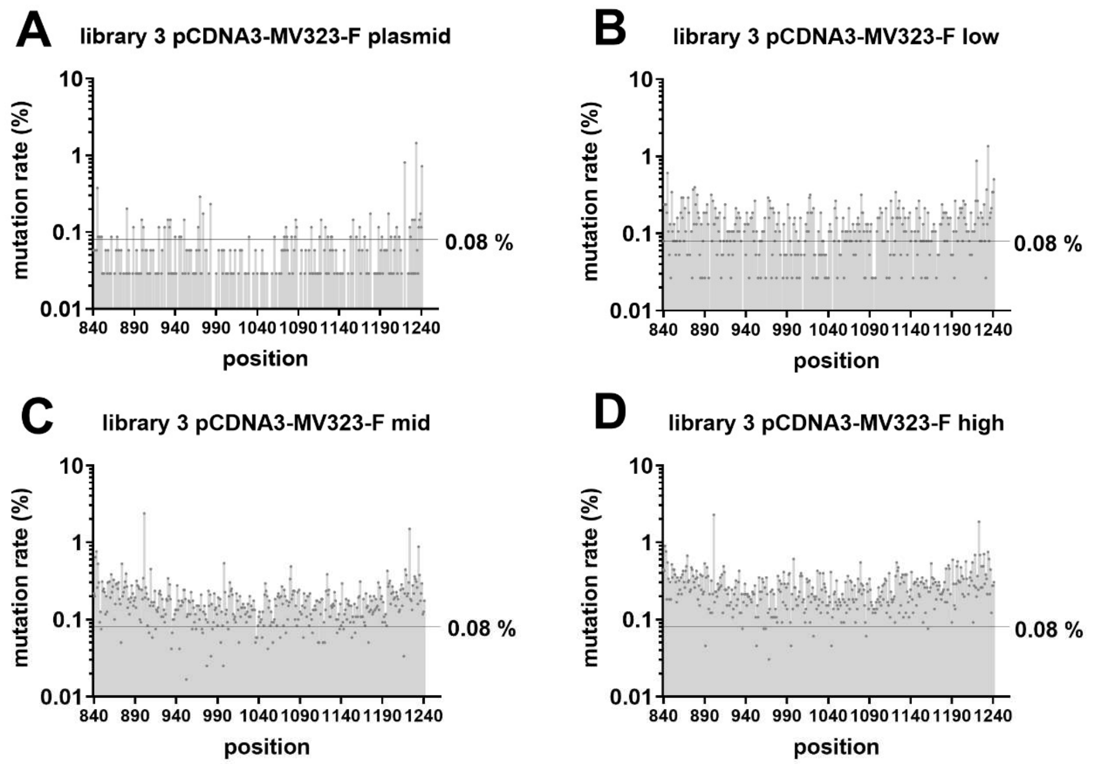


**E**

| shuttle libraries | library 3 | library 3 | library 3 | library 3 |
|---|---|---|---|---|
| mutation setting | plasmid DNA | low | medium | high |
| average mutation rate (%) | 0.04% | 0.12% | 0.18% | 0.27% |
| Ts/Tv | 0.4 | 0.81 | 1.03 | 0.81 |
| coverage (%) | 15.7% | 52.2% | 89.8% | 97.3% |
| number of bacterial colonies retrieved | - | 1.08x10^5 | 7.47x10^4 | 3.15x10^4 |

**Fig. S1. Optimization of error-prone PCR conditions to generate saturation mutagenesis library of MeV-F**. The background mutation rate of library 3 (MeV-F ORF, nt 840-1241) amplified from the shuttle plasmid DNA by high fidelity DNA polymerase represents the intrinsic error rate of Illumina sequencing under our conditions (A). The mutation rates of library 3 generated with the low (B), medium (C), and high (D) error-prone PCR conditions (mutational setting) as per manufacturer's recommendations (Genemorph II random mutagenesis kit, Agilent) show the expected increase. Position on the x-axis indicates the nucleotide position counted from initiation codon of F ORF (ATG, nt 1-3).

The mutation profile of library 3 for the aforementioned conditions are summarized in (E). 'Ts/Tv' stands for transversion / transition rate. Nucleotide positions with a mutation rate of $\geq$ 0.08% (2X 'plasmid DNA' background rate of 0.04%) are defined as having real mutations. The % coverage across library 3 for each condition was then calculated by using 0.08% as a cut-off for determining whether that nucleotide position contained a genuine (selectable) mutation.

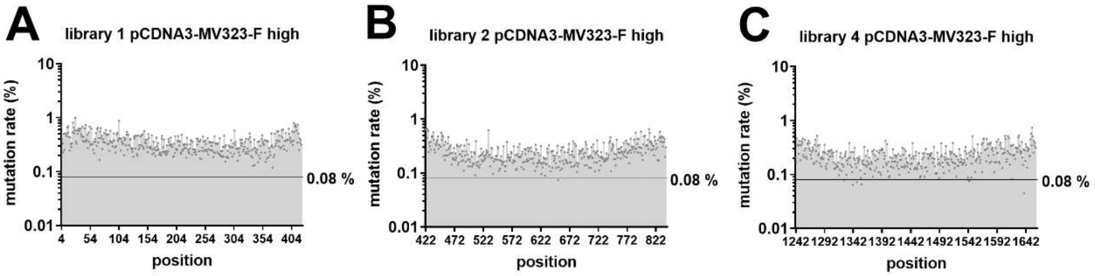


D

| shuttle libraries | library 1 | library 2 | library 4 |
|---|---|---|---|
| mutation setting | high | high | high |
| average mutation rate (%) | 0.33% | 0.27% | 0.22% |
| Ts/Tv | 0.99 | 0.88 | 0.97 |
| coverage (%) | 100% | 100% | 98.3% |
| number of bacterial colonies retrieved | 1.64x10^5 | 1.50x10^5 | 2.62x10^4 |

**Fig. S2. Mutation profile of shuttle plasmid libraries in library 1, 2, and 4.** (A) to (C) show the mutation rates of shuttle plasmid library 1, 2, and 4 with high mutation settings. Position means the nucleotide position counted from initiation codon of F open reading frame. (D) shows the summarized mutation profile of shuttle plasmid library 1, 2, and 4. ‘Ts/Tv’ stands for transversion / transition rate. Nucleotides with mutation rate of >= 0.08% is defined as real mutation, then coverages were calculated by using 0.08% as a lower detection limit of mutation.

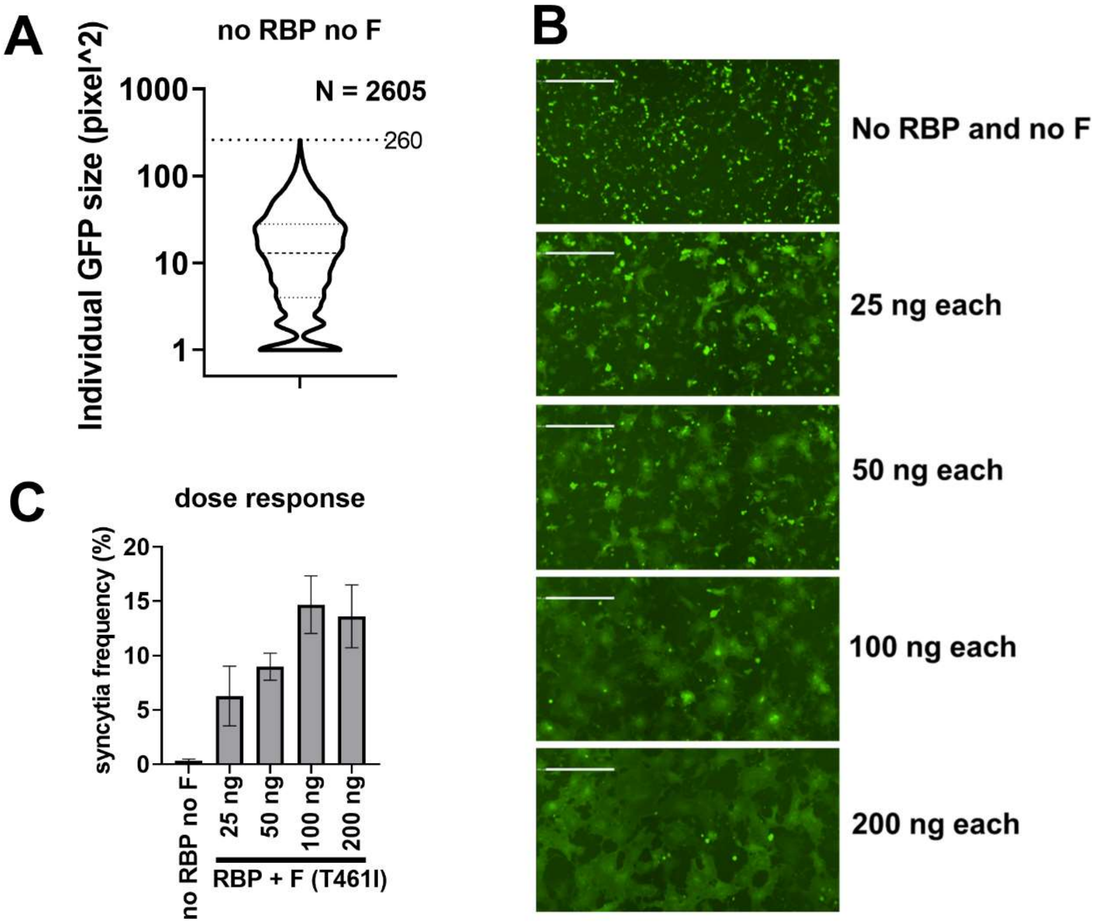


**Fig. S3. Validation of image based fusion assay.** MeV-RBP, MeV-F (T461I), and Lifeact-GFP expressing plasmids were transfected to BSR-T7 cells. Syncytia formation was visualized by Celigo imaging cytometer and quantified by ImageJ. (A) shows the distribution of individual GFP size (without fusion) from no RBP and no F transfection of replicate 1 experiment. Dot lines show quartiles (75% and 25%) and median (50%) of population. Syncytia frequency (details are in the method) was used as value showing fusogenicity. Syncytia frequency elevated with the increasing amount of RBP and F-T461I transfection. Value is shown as average of 5 independent experiments with error-bar indicating standard deviation. (B) shows representative images from fusion assay. (C) shows correlation between the fusion activity and the amount RBP plus F-T461I transfection. Images in (B) are taken by Celigo Imaging Cytometer (Nexcelom), and they are computational composite of several identical fields of view taken in each well. White bar in the picture is equal to 500 micrometer.

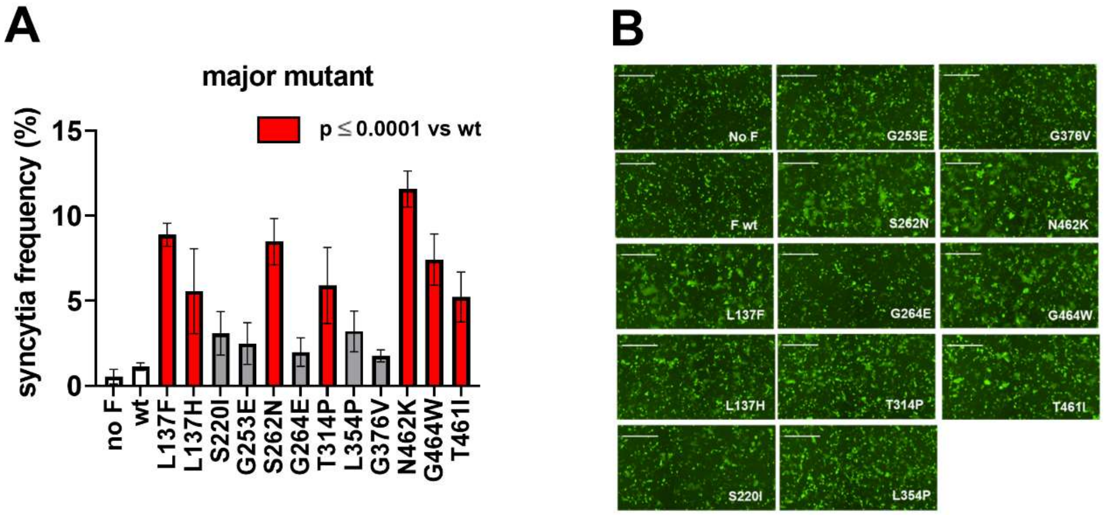


**Fig. S4. Image based fusion assay for distinguishing the hyper fusion mutants.** MeV-RBP 25ng, MeV-F 25ng and Lifeact-GFP expressing plasmids were transfected to BSR-T7 cells to distinguish the fusogenicity among hyper fusion mutants. Fusion was evaluated 30 hours after transfection (A). Values are shown as average of 5 independent experiments with error-bar indicating standard deviation. Dunnet's multiple comparison test was used for the detection of statistical significant difference above wild type transfection. The representative images in the fusion assay were shown (B). All images shown here are taken by Celigo Imaging Cytometer (Nexcelom).

**Table S1. Mutation profile in genome plasmid libraries and rescue efficiency in the screening experiments.** The table shows the mutation profile of genome plasmid libraries, and rescue efficiency in the screening experiments. 'Ts/Tv' stands for transversion / transition rate. Nucleotides with mutation rate of ≥0.08% is defined as real mutation, then coverages were calculated by using 0.08% as a lower detection limit of mutation. The number of GFP positive cells was counted at day2 post rescue trial and used as a measure to evaluate rescue efficiency.

| genome libraries | library 1 | library 2 | library 3 | library 4 |
|---|---|---|---|---|
| average mutation rate (%) | 0.35% | 0.25% | 0.26% | 0.14% |
| Ts/Tv | 1.02 | 0.88 | 0.84 | 0.98 |
| coverage (%) | 100% | 99.8% | 99.0% | 85.0% |
| number of bacterial colonies retrieved | 3.53x10^4 | 8.55x10^4 | 3.02x10^4 | 2.32x10^4 |
| rescue trial | library 1 | library 2 | library 3 | library 4 |
| Average GFP positive cells / well | 8.2x10^4 | 6.3x10^4 | 8.7x10^4 | 8.1x10^4 |
| S.D. of GFP positive cells | 1.8x10^4 | 2.4x10^4 | 2.5x10^4 | 2.5x10^4 |

**Table S2. Top hit list in the screening experiments.** The most predominant mutants in each library are ranked by their mutation rate (%) and listed along with the indicated information about that mutant. We conducted three independent rescue and virus outgrowth experiments for each of the four libraries (lib1, 2, 3, and 4). Synonymous mutations that appeared in the top hit list are almost always associated with non-synonymous mutations, suggesting that these non-synonymous passenger mutations are real, but neutral. Highlighted in magenta are the top two non-synonymous mutations from each library that we selected for functional validation. Nt; nucleotide. AA; amino acid. ON; original nucleotide (original plasmid sequence). MN; mutated nucleotide. Sub; amino acid substitution.

| | lib1 | | | | | | lib1 | | | | | | lib1 | | | | | |
|---|---|---|---|---|---|---|---|---|---|---|---|---|---|---|---|---|---|---|
| | replicate1 | | | | | | replicate2 | | | | | | replicate3 | | | | | |
| rank | Nt | AA | % | ON | MN | Sub | Nt | AA | % | ON | MN | Sub | Nt | AA | % | ON | MN | Sub |
| 1 | 409 | 137 | 64.1% | C | T | L137F | 51 | 17 | 10.6% | T | A | syn | 410 | 137 | 57.8% | T | A | L137H |
| 2 | 351 | 117 | 12.2% | C | A | syn | 287 | 96 | 10.3% | A | C | Q96P | 409 | 137 | 24.1% | C | T | L137F |
| 3 | 410 | 137 | 6.6% | T | A | L137H | 394 | 132 | 5.7% | A | T | T132S | 4 | 2 | 7.5% | G | T | G2C |
| 4 | 164 | 55 | 5.2% | T | A | V55D | 410 | 137 | 5.3% | T | A | L137H | 219 | 73 | 7.1% | C | G | I73M |
| 5 | 117 | 39 | 5.2% | A | G | syn | 101 | 34 | 4.5% | G | A | G34E | 51 | 17 | 2.4% | T | A | syn |
| 6 | 287 | 96 | 4.4% | A | T | Q96L | 270 | 90 | 4.3% | A | G | syn | 287 | 96 | 1.9% | A | C | Q96P |
| 7 | 87 | 29 | 4.0% | T | C | syn | 6 | 2 | 4.3% | T | C | syn | 416 | 139 | 1.8% | A | T | Q139L |
| 8 | | | | | | | 31 | 11 | 4.3% | T | C | F11L | 217 | 73 | 1.8% | A | T | I73F |
| 9 | | | | | | | 106 | 36 | 4.2% | G | A | V36I | 72 | 24 | 1.8% | A | T | Q24H |
| note | Most 351 syn are on the same read of 409 mut | | | | | | 51 syn and 287 mut are on the same reads | | | | | | 4 mut are on the same read of 410 or 409 mut. | | | | | |
| | lib2 | | | | | | lib2 | | | | | | lib2 | | | | | |
| | replicate1 | | | | | | replicate2 | | | | | | replicate3 | | | | | |
| rank | Nt | AA | % | ON | MN | Sub | Nt | AA | % | ON | MN | Sub | Nt | AA | % | ON | MN | Sub |
| 1 | 655 | 219 | 35.7% | C | A | P219T | 791 | 264 | 26.2% | G | A | G264E | 659 | 220 | 16.7% | G | T | S220I |
| 2 | 810 | 270 | 27.7% | T | A | syn | 837 | 279 | 26.1% | T | A | syn | 460 | 154 | 16.5% | C | T | syn |
| 3 | 785 | 262 | 27.3% | G | A | S262N | 460 | 154 | 17.4% | C | T | syn | 779 | 260 | 15.0% | T | C | L260S |
| 4 | 424 | 142 | 6.7% | C | A | L142M | 758 | 253 | 17.2% | G | A | G253E | 751 | 251 | 8.9% | T | A | K251N |
| 5 | 679 | 227 | 6.4% | G | C | A227P | 793 | 265 | 4.1% | A | G | I265V | 677 | 226 | 8.4% | C | T | S226F |
| 6 | 517 | 173 | 5.6% | T | C | syn | 663 | 221 | 3.9% | A | C | L221F | 651 | 217 | 6.4% | T | A | F271L |
| 7 | 719 | 240 | 5.4% | G | A | G240E | 795 | 265 | 3.0% | A | T/G | I265I/M | 776 | 259 | 5.7% | T | C | I259T |
| 8 | 651 | 217 | 4.3% | T | G | F217L | | | | | | | 550 | 184 | 4.5% | A | G | N184D |
| 9 | | | | | | | | | | | | | 717 | 239 | 4.5% | A | T | syn |
| note | Most 810 syn and 785 mut are on the same reads | | | | | | 837 and 460 syn are either on 791 or 758 mut. | | | | | | 659 mut and 460 syn are moving together | | | | | |
| | lib3 | | | | | | lib3 | | | | | | lib3 | | | | | |
| | replicate1 | | | | | | replicate2 | | | | | | replicate3 | | | | | |
| rank | Nt | AA | % | ON | MN | Sub | Nt | AA | % | ON | MN | Sub | Nt | AA | % | ON | MN | Sub |
| 1 | 940 | 314 | 58.9% | A | C | T314P | 1127 | 376 | 54.8% | G | T | G376V | 1125 | 375 | 19.6% | T | A | F375L |
| 2 | 1127 | 376 | 16.2% | G | T | G376V | 850 | 284 | 40.6% | G | A | A284T | 1061 | 354 | 11.8% | T | C | L354P |
| 3 | 851 | 284 | 15.9% | C | T | A284V | 889 | 297 | 21.8% | C | T | H297Y | 851 | 284 | 8.6% | C | T | A284V |
| 4 | 1061 | 354 | 6.4% | T | A | L354Q | 851 | 284 | 14.0% | C | T | A284V | 850 | 284 | 8.1% | G | A | A284T |
| 5 | 850 | 284 | 4.3% | G | A | A284T | 1213 | 405 | 5.4% | A | T | N405Y | 1239 | 413 | 7.0% | A | G | syn |
| 6 | 907 | 303 | 4.3% | T | A | S303T | 842 | 281 | 2.9% | T | C | L281P | 1187 | 396 | 6.8% | A | G | K396R |
| 7 | 1141 | 381 | 2.7% | T | A | L381I | 843 | 281 | 2.1% | C | T | syn | 1008 | 336 | 6.4% | C | T | syn |
| 8 | 1089 | 363 | 2.5% | C | G | syn | 846 | 282 | 1.7% | T | C | syn | 907 | 303 | 4.6% | T | A | S303T |
| note | 1127 and 851 mut are manly on single read | | | | | | mutants are overlapped each other | | | | | | | | | | | |
| | lib4 | | | | | | lib4 | | | | | | lib4 | | | | | |
| | replicate1 | | | | | | replicate2 | | | | | | replicate3 | | | | | |
| rank | Nt | AA | % | ON | MN | Sub | Nt | AA | % | ON | MN | Sub | Nt | AA | % | ON | MN | Sub |
| 1 | 1300 | 434 | 0.79% | A | G | S434G | 1386 | 462 | 7.3% | T | A | N462K | 1390 | 464 | 1.75% | G | T | G464W |
| 2 | 1629 | 543 | 0.62% | A | G | syn | 1393 | 465 | 1.4% | A | T | N465Y | 1298 | 433 | 0.84% | G | A | G433E |
| 3 | 1613 | 538 | 0.53% | A | G | D538G | 1442 | 481 | 1.2% | A | G | D481G | 1386 | 462 | 0.58% | T | G | N462K |
| 4 | 1607 | 536 | 0.51% | A | G | K536R | 1568 | 523 | 0.97% | A | G | K523R | 1394 | 465 | 0.48% | A | G | N465S |
| 5 | 1309 | 437 | 0.47% | T | C | Y437H | 1476 | 492 | 0.96% | C | G | S492R | 1277 | 426 | 0.43% | A | G | N426S |
| 6 | 1575 | 525 | 0.45% | A | G | syn | 1591 | 531 | 0.96% | T | C | S531P | 1624 | 542 | 0.41% | A | G | T542A |
| 7 | | | | | | | 1497 | 499 | 0.93% | G | A | syn | 1453 | 485 | 0.41% | A | G | R485G |
| 8 | | | | | | | 1613 | 538 | 0.91% | A | G | D538G | 1464 | 488 | 0.38% | A | G | syn |
| 9 | | | | | | | | | | | | | 1552 | 518 | 0.38% | A | G | R518G |

**Table S2. Top hit list in preliminary optimization screening experiment using library 3.** Position and mutation rate in passage 1 and passage 2 are shown. Nt; nucleotide. AA; amino acid. ON; original nucleotide (original plasmid sequence). MN; mutated nucleotide. Sub; amino acid substitution.

| | lib3 (preliminary experiment) | | | | | | | | | | | |
|---|---|---|---|---|---|---|---|---|---|---|---|---|
| | Passage 1 | | | | | | Passage 2 | | | | | |
| rank | nt | aa | % | ON | MN | sub | nt | aa | % | ON | MN | sub |
| 1 | 1127 | 376 | 36.0% | G | T | G376V | 1127 | 376 | 77.8% | G | T | G376V |
| 2 | 1123 | 375 | 11.8% | T | G | F375V | 1179 | 393 | 17.7% | C | G | I393M |
| 3 | 850 | 284 | 2.1% | G | A | A284T | 882 | 294 | 17.4% | G | A | syn |
| 4 | 875 | 292 | 1.3% | A | T | K292M | 889 | 297 | 17.4% | C | T | H297Y |
| 5 | | | | | | | 907 | 303 | 1.5% | T | A | S303T |
| note | | | | | | | 882, 889, and 1179 are on the same reads. | | | | | |

**Table S4. Summary list of mutations in the MeV-F protein that confer a hyperfusogenic SSPE/MIBE phenotype.** Previously described mutations are indicated in black[1–3] while novel mutations selected by our screen and functionally characterized in the current study are in red. The mutants are sorted by position in each of the four libraries (lib1 - lib 4) that make up our screen of the MeV-F protein. Grey highlighted rows indicate SSPE mutations that must occur together in order to confer hyperfusogenicity [2]. Yellow highlighted rows indicate different mutations of the same residue.

| lib1 | mean RMF[1] | input (%)[2] |
|---|---|---|
| I87T | 0.7 | 0.12 |
| M94V | 0.95 | 0.12 |
| Q96P | 35.7 | 0.04 |
| S103I | 1.4 | 0.09 |
| L137H | 52.3 | 0.36 |
| L137F | 53.3 | 0.43 |

| lib2 | mean RMF[1] | input (%)[2] |
|---|---|---|
| G168R | 0.58 | 0.11 |
| E170G | 1.7 | 0.14 |
| P219T | 33.6 | 0.08 |
| S220I | 33.5 | 0.05 |
| G253E | 23.4 | 0.14 |
| L260S | 30.5 | 0.12 |
| S262R[3a] | < 0.1 | 0.14 |
| S262N[3b] | 25.7 | 0.12 |
| G264E | 33.6 | 0.14 |

| lib3 | mean RMF[1] | input (%)[2] |
|---|---|---|
| A284T | 40.9 | 0.26 |
| H297Y | 15.9 | 0.13 |
| T314P | 33.8 | 0.07 |
| L354M[3a] | < 0.1 | 0.11 |
| L354P[3b] | 23.6 | 0.08 |
| G376V | 57.0 | 0.18 |

| lib4 | mean RMF[1] | input (%)[2] |
|---|---|---|
| A440P | < 0.1 | < 0.01 |
| L454W | < 0.1 | < 0.01 |
| T461I[3a] | < 0.1 | 0.06 |
| T461A[3b] | 20.6[3c] | 0.32 |
| N462K[4] | 51.7 | 0.45 |
| N462S[4] | 17.9 | 0.25 |
| G464W[3b] | 37.5[3c] | 1.67 |
| G464E[3a] | < 0.1 | 0.02 |
| N465S[3b] | 23.9[3c] | 0.31 |
| N465K[3a] | < 0.1 | 0.01 |
| R520C | < 0.1 | 0.09 |
| L550P | < 0.1 | 0.13 |
| F long tail | | |
| F delta-30 | | |

[1] Relative Mutation Frequency (RMF) calculated as described in Fig. 4E.
[2] The prevalence of mutations in the input library that give rise to that particular amino acid change indicated for that hyperfusogenic mutant.
[3] (a) Hyperfusogenic mutant described in the literature that was not recovered in our library screen but a different mutation at the same residue (b) was positively selected as indicated by its high mean RMF value (c).
[4] Distinct mutations at the same residue reported to result in a hyperfusogenic SSPE phenotype that were also positively selected in our screen.

**Table S5. List of primer set in this study.**

| | primer name | sequence |
|---|---|---|
| primer set for error-prone PCR (insert) | | |
| for library 1 | library 1 ins-f | ACTCATCCAGTGTCCATCATG |
| | library 1 ins-r | GATGGCTTGAGAGTTCAGCA |
| for library 2 | library 2 ins-f | CATTGCACTTCACCAGTCCA |
| | library 2 ins-r | GTCGGATAGGCTATACTGAGT |
| for library 3 | library 3 ins-f | GACACAGAGTCCTACTTCATTGT |
| | library 3 ins-r | GCAGTGATCGGCAGCAATG |
| for library 4 | library 4 ins-f | GACCCTGACAAGATCCTAACATA |
| | library 4 ins-r | TGTTTCAAGAGTTGTAGAGGATCA |
| pCDNA3-MV-F backbone primer for shuttle library | | |
| for library 1 | library 1 back-f | GCTGAACTCTCAAGCCATCGAC |
| | library 1 back-r | TGATGGACACTGGATGAGTCTTGA |
| for library 2 | library 2 back-f | TCAGTATAGCCTATCCGACGCTG |
| | library 2 back-r | GGACTGGTGAAGTGCAATGCC |
| for library 3 | library 3 back-f | CATTGCTGCCGATCACTGCC |
| | library 3 back-r | GTAGGACTCTGTGTCGACGTGAG |
| for library 4 | library 4 back-f | CTCTACAACTCTTGAAACACAGATTTCCC |
| | library 4 back-r | TTAGGATCTTGTCAGGGTCTTGATTAATG |
| 1st nested PCR (RT-PCR) | | |
| nested RT-PCR 1st | nested 1st-f | ACTCATCCAGTGTCCATCATG |
| | nested 1st-r | TGTTTCAAGAGTTGTAGAGGATCA |
| Amplicon PCR primer (for 2nd nested PCR) | | |
| for library 1 | nested 2nd lib1-f | CCAGTGTCCATCATG |
| | nested 2nd lib1-r | CTTGAGAGTTCAGCA |
| for library 2 | nested 2nd lib2-f | CACTTCACCAGTCCA |
| | nested 2nd lib2-r | TAGGCTATACTGAGT |
| for library 3 | nested 2nd lib3-f | GACACAGAGTCCTACTTCATTGT |
| | nested 2nd lib3-r | GCAGTGATCGGCAGCAATG |
| for library 4 | nested 2nd lib4-f | GACAAGATCCTAACATA |
| | nested 2nd lib4-r | AAGAGTTGTAGAGGATCA |
| genome specific RT-qPCR primer | | |
| for genome RT | MV genome-f | accaaacaaagttgggtaagga |
| qPCR primer | EGFP-f | acgtaaacggccacaagttc |
| | EGFP-r | aagtcgtgctgctycatgtg |
| mutants cloning primer | | |
| Q96P | Q96P-f | AATGACCCCGAATATAAGACCGGTTCAGA |
| | Q96P-r | ATATTCGGGGTCATTGCATTAAGTGCATCTCT |
| T132S | T132S-f | TCAGATATCAGCCGGCATTGCACTTCACCA |
| | T132S-r | CCGGCTGATATCTGAGCAGCTGTGGCAAC |
| L137F | L137F-f | TTGCATTTCACCAGTCCATGCTGAACT |
| | L137F-r | ACTGGTGAAATGCAATGCCGGCTGTTATCTG |
| L137H | L137H-f | ATTGCACATCACCAGTCCATGCTGAAC |
| | L137H-r | CTGGTGATGTGCAATGCCGGCTGTTATCTG |
| P219T | P219T-f | ATTTGGCACCAGCTTACGGGACCCCATAT |
| | P219T-r | AAGCTGGTGCCAAATAATGACAGGATTTCTG |
| S220I | S220I-f | TGGCCCCATCTTACGGGACCCCATATCT |
| | S220I-r | CGTAAGATGGGGCCAAATAATGACAGGATT |
| G253E | G253E-f | ATACAGTGAAGGTGATTTACTGGGCATCTTAGA |
| | G253E-r | TCACCTTCACTGTATCCGAGCTTTTCTAAT |
| L260S | L260S-f | GGGCATCTCAGAGAGCAGAGGAATAAAGG |
| | L260S-r | CTCTCTGAGATGCCCAGTAAATCACCTCCA |
| S262N | S262N-f | CTTAGAGAACAGAGGAATAAAGGCCCGGAT |

| | | |
|---|---|---|
| | S262N-r | CCTCTGTTCTCTAAGATGCCCAGTAAATCACCT |
| G264E | G264E-f | GAGCAGAGAAATAAAGGCCCGGATAACTCA |
| | G264E-r | TTTATTTCTCTGCTCTCTAAGATGCCCAGTAAA |
| A284T | A284T-f | TCAGTATAACCTATCCGACGCTGTCCGAG |
| | A284T-r | GATAGGTTATACTGAGTACAATGAAGTAGGACTC |
| H297Y | H297Y-f | TGTCTACCGGCTAGAGGGGGTCTC |
| | H297Y-r | TCTAGCCGGTAGACAATCACCCCCT |
| T314P | T314P-f | GGTATACCCCTGTGCCCAAGTATGTTGCA |
| | T314P-r | GCACAGGGGTATACCACTCTTGAGAGCCT |
| L354P | L354P-f | GAGTCCTCCGCTCCAAGAATGCCTCC |
| | L354P-r | TGGAGCGGAGGACTCATCGGGTACAAG |
| F375L | F375L-f | GGGTCTTTAGGGAACCGGTTCATTTTATC |
| | F375L-r | GTTCCCTAAAGACCCGGATACGAGTGTA |
| F375V | F375V-f | GGTCTGTTGGGAACCGGTTCA |
| | F375V-r | GGTTCCCAACAGACCCGGATA |
| G376V | G376-f | TCTTTTGTGAACCGGTTCATTTTATC |
| | G376-r | CCGGTTCACAAAAGACCCGGA |
| I393M | F393 f | ATCAATGCTCTGCAAGTGTTACACAA |
| | F393 r | TTGCAGAGCATTGATGCACAATTGGCT |
| G433E | G433E-f | CCAAGTCGAGAGCAGGAGGTATCCGGA |
| | G433E-r | CTGCTCTCGACTTGGATGGTCACACCGTT |
| S434G | S434G-f | AAGTCGGGGGCAGGAGGTATCCGGACG |
| | S434G-r | TCCTGCCCCCGACTTGGATGGTCAC |
| T461I | T461I-f | GTAGGGATAAATCTGGGGAATGCAATTG |
| | T461I-r | CCAGATTTATCCCTACGTCCAACCTCTC |
| G464W | G464W-f | CAAATCTGTGGAATGCAATTGCTAAGTTG |
| | G464W-r | CATTCCACAGATTTGTCCCTACGTCCAAC |
| N465Y | N465Y-f | ATCTGGGGTATGCAATTGCTAAGTTGGA |
| | N465Y-r | TTGCATACCCCAGATTTGTCCCTACGTC |
| D481G | D481G-f | TCATCGGGCCAGATATTGAGGAGTATGAAAGGT |
| | D481G-r | TATCTGGCCCGATGACTCCAACAATTCCTTG |
| D538G | D538G-f | AAAGCCTGGTCTTACAGGGACATCAAAA |
| | D538G-r | GTAAGACCAGGCTTTAGGCCTGGTCTTG |
